# Gene module-trait network analysis uncovers cell type specific systems and genes relevant to Alzheimer’s Disease

**DOI:** 10.1101/2025.01.31.635970

**Authors:** Katia de Paiva Lopes, Ricardo A. Vialle, Gilad Green, Masashi Fujita, Chris Gaiteri, Vilas Menon, Julie A. Schneider, Yanling Wang, Philip L. De Jager, Naomi Habib, Shinya Tasaki, David A. Bennett

## Abstract

Alzheimer’s Disease (AD) is marked by the accumulation of pathology, neuronal loss, and gliosis and frequently accompanied by cognitive decline. Understanding brain cell interactions is key to identifying new therapeutic targets to slow its progression. Here, we used systems biology methods to analyze single-nucleus RNA sequencing (snRNASeq) data generated from dorsolateral prefrontal cortex (DLPFC) tissues of 424 participants in the Religious Orders Study or the Rush Memory and Aging Project (ROSMAP). We identified modules of co-regulated genes in seven major cell types, assigned them to coherent cellular processes, and assessed which modules were associated with AD traits such as cognitive decline, tangle density, and amyloid-β deposition. Coexpression network structure was conserved in the majority of modules across cell types, but we also found distinct communities with altered connectivity, especially when compared to bulk RNASeq, suggesting cell-specific gene co-regulation. These coexpression modules can also capture signatures of cell subpopulations and be influenced by cell proportions. Using a Bayesian network framework, we modeled the direction of relationships between the modules and AD progression. We highlight two key modules, a microglia module (mic_M46), associated with tangles; and an astrocyte module (ast_M19), associated with cognitive decline. Our work provides cell-specific molecular networks modeling the molecular events leading to AD.

## Introduction

Alzheimer’s Disease (AD), the most common cause of dementia, is characterized by the accumulation of neuritic plaques and neurofibrillary tangles, accompanied by neuronal loss and gliosis. The most common clinical consequence is loss of cognition, especially episodic memory, with variable symptoms across patients (Knopman et al. 2021; Jack et al. 2018). Despite decades of research, no therapies exist to meaningfully treat or prevent AD. The complexity and heterogeneity of AD still presents a significant challenge to understanding its pathophysiology, necessitating interdisciplinary approaches to tackle the problem from different angles.

Recent advances in genetic studies have implicated pathways in distinct cell-types involved with the disease development (Bellenguez et al. 2022; Yang et al. 2023; Yap et al. 2024). Supporting this, single-cell and single-nucleus RNA sequencing (snRNASeq) profiling from brain tissues, have shown that AD involves a complex interplay of every major brain cell type (Grubman et al. 2019). For instance, there are specific subpopulations of neurons vulnerable to AD while other groups of cells may be resilient to AD pathology. Glial cell populations, namely astrocyte, oligodendrocytes, and microglial cells, are highly dynamic and respond differently to distinct environmental signals in the damaged brain (Allen and Lyons 2018; Gazestani et al. 2023; Murdock and Tsai 2023). Therefore, a deep understanding of the molecular pathways in specific cell types and their responses to AD is crucial for developing more targeted and effective therapies.

We previously showed that genetic variation and gene expression exhibit cell type and subtype-specific associations in the brain. (Fujita et al. 2024; Green et al. 2024). Here, we describe the module-trait network (MTN) approach for finding and modeling cell-specific gene regulatory networks of the aging brain. We analyzed snRNASeq from human dorsolateral prefrontal cortex (DLPFC) tissues of 424 participants from the Religious Orders Study or the Rush Memory and Aging Project (ROSMAP). By constructing single-nucleus coexpression networks for each of the major cell types of the cortex we uncovered the intricate relationships between genes, their functions, and their associations with AD traits. We then conducted detailed module characterization, including comparison of coexpression network structure between cell types, gene ontology (GO) and pathway enrichment analyses, associations with subpopulations of cells, and correlations with various cognitive performance tests and brain pathologies. This comprehensive set of data-driven, cell-specific molecular modules was then integrated into a Bayesian network framework to model the cascade of multi-cellular molecular events leading to AD development (**Supplementary** Figure 1).

## Results

### Samples description

Transcriptomic data originate from two longitudinal clinical-pathologic cohort studies of aging and dementia, the Religious Orders Study and Rush Memory and Aging Project (ROSMAP). Participants are older adults who enroll without known dementia and agree to annual clinical evaluations and brain donation. To date, ROSMAP has enrolled over 4,000 participants and there have been more than 2,000 brain autopsies. Both studies were approved by an Institutional Review Board of Rush University Medical Center. All participants signed informed and repository consents and an Anatomic Gift Act. The snRNASeq transcriptomics were profiled from postmortem DLPFC brain tissues of 424 individuals while bulk RNASeq from the DLPFC were available for 1,210 participants (**Supplementary Table 1**). The participants enrolled at a mean age of 80.8 (SD: 7.0) years, and died at an average age of 89.5 (SD: 6.6), with more than two thirds being female (∼70%). At death, 35% had no cognitive impairment (NCI), 25% had mild cognitive impairment (MCI), and the remaining 40% had dementia. At autopsy, 64% had a pathologic AD. TDP-43 pathology extending beyond the amygdala (limbic-predominant age-related TDP-43 encephalopathy - LATE) was detected in 30% of the brains (**Supplementary Table 2**). The characteristics were similar for individuals with snRNASeq data (**Supplementary Table 3**).

### The single-nucleus coexpression networks of the aging human brain

We applied the MTN approach to summarize the transcriptome-wide information into coherent modules of genes whose final goal is to prioritize specific systems and inform on hypotheses for further experiments. The MTN approach consists of the following steps (C. Gaiteri et al. 2014): 1) Construct coexpression networks; 2) Identify groups of coexpressed genes that represent molecular systems; and 3) Model directional relationships between the modules and AD traits using Bayesian networks. We obtained the processed and annotated snRNASeq data and created participant-level normalized pseudo-bulk matrices for each cell type group annotated for the DLPFC (syn31512863, **Figure 1 a-c** and **Supplementary** Figure 1-2, **Methods**) (Fujita et al. 2024; Green et al. 2024). We then applied Speakeasy (Chris Gaiteri et al. 2015) to identify single-nucleus coexpressed networks of the human DLPFC. We found 193 modules with at least 30 genes each, across the major cell types of the DLPFC, comprising 26 modules for astrocytes, 26 for endothelial cells, 29 for excitatory neurons, 24 for inhibitory neurons, 30 for microglial cells, 30 for oligodendrocytes, and 28 for oligodendrocyte precursor cells (OPCs) (**Supplementary** Figure 3). The genes assigned to each module are available in **Supplementary Tables 4-10**. All the coexpression networks followed a scale-free topology where a few hub genes present a large number of connections, as expected in real-world biological data (**Supplementary** Figure 4).

**Fig. 1.**
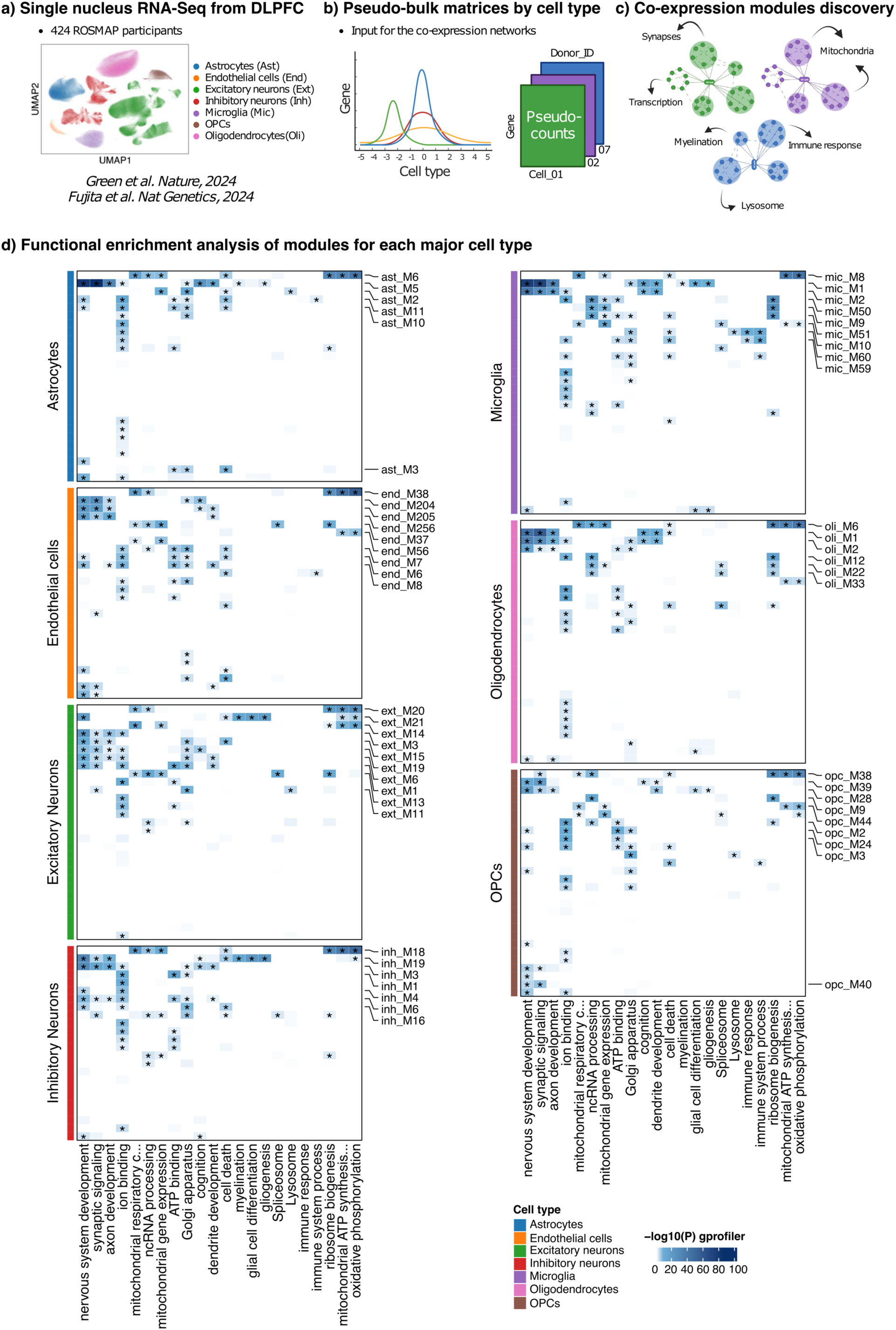
The single-nucleus coexpression networks of the human cortex. **a)** The ROSMAP snRNASeq data used for the networks covers 1.6 M cells, 7 major cell-type groups, and 96 cell subpopulations (Fujita et al. 2024; Green et al. 2024). **b)** Pseudo-bulk matrices were created and used as input for the coexpression networks. **c)** Coexpression modules were identified using Speakeasy clustering algorithm (Chris Gaiteri et al. 2015). **d)** Heatmap showing the Functional enrichment analysis for each module for selected Gene Ontology (GO) terms. Modules are organized by cell type, with each module labeled according to its corresponding cell type and module number (e.g., ast_M6 for astrocyte module 6). The significance of enrichment is represented by a gradient of -log_10_(*P*-value), where darker colors indicate stronger enrichment. The *P*-values were adjusted for multiple testing from gProfiler’s “g_SCS” function. Asterisks mark the modules with adj *P*-value < 0.05.

Our coexpression modules captured a variety of biological systems within each cell type, as demonstrated by the functional enrichment analysis (**Figure 1d**). As expected, ubiquitous eukaryotic cell functions were found enriched in modules from all cell types, including terms such as cell death, oxidative phosphorylation, ATP synthesis, mitochondrial respiration, and ribosomal biogenesis. Terms related with brain tissues, such as nervous system development, and synaptic organization, were also shared across all cell types. By contrast, other well-known cell-type-specific functions highlighted the specialized roles of modules in each cell type. For instance, immune-related terms were enriched mostly in modules derived from microglial cells, while other specific functions like gliogenesis and lysosome were found in modules from a subset of cell types including microglia, astrocytes, OPCs, and neurons. Overall, this analysis highlights the balance between biological processes shared in all cortical cells and specialized functions critical for individual cell types in maintaining cortical structure and function.

### Module similarity across cell types and bulk RNASeq

To assess the overall similarity of modules in each cell type, we estimated the distances between modules based on their module preservation and normalized mutual information results, and for replication, we compared these with modules derived from bulk RNASeq. Such a comparison has the potential to highlight biological signals that are lost at the bulk level, particularly in less prevalent cell types or specific cell subpopulations. We used two bulk RNASeq datasets: one previously published by Mostafavi et al. (2018), which included 478 samples, and our updated version with additional samples totaling 1,210 participants (**Supplementary Tables 11-13** and **Supplementary** Figures 5-8). Principal component analysis (PCA) of the distances revealed consistent clustering patterns with both module preservation and normalized mutual information metrics. Specifically, excitatory and inhibitory neuron modules grouped together, followed by clusters of oligodendrocyte-, astrocyte-, and OPC-related modules, with microglial and endothelial modules clustering together. Also, all single-nucleus modules were distant from those derived from bulk RNASeq data (**Figure 2a – b**) and both module preservation and normalized mutual information results showed strong concordance in differentiating these patterns (**Figure 2c**). Every cell type tested had gene modules preserved in the bulk dataset, with moderate preservation defined as Zsummary ≥ 2 and high preservation as Zsummary ≥ 10. A total of 56 modules were not preserved in bulk RNASeq (Zsummary < 2), with numbers ranging from 6 to 11 specific modules depending on the cell type population (**Figure 2d**). For example, nine microglia modules (mic_M16, mic_M34, mic_M45, mic_M46, mic_M50, mic_M52, mic_M55, mic_M64 and mic_M65) while 11 excitatory neuron modules (ext_M2, ext_M4, ext_M5, ext_M7, ext_M10, ext_M23, ext_M26, ext_M27, ext_M28, ext_M29 and ext_M30) were not preserved in the bulk coexpression networks (**Figure 2e–f**). Pairwise preservation metrics, the complete module preservation and normalized mutual information results are available on the **Supplementary Tables 14-16** and **Supplementary** Figure 9.

**Fig. 2.**
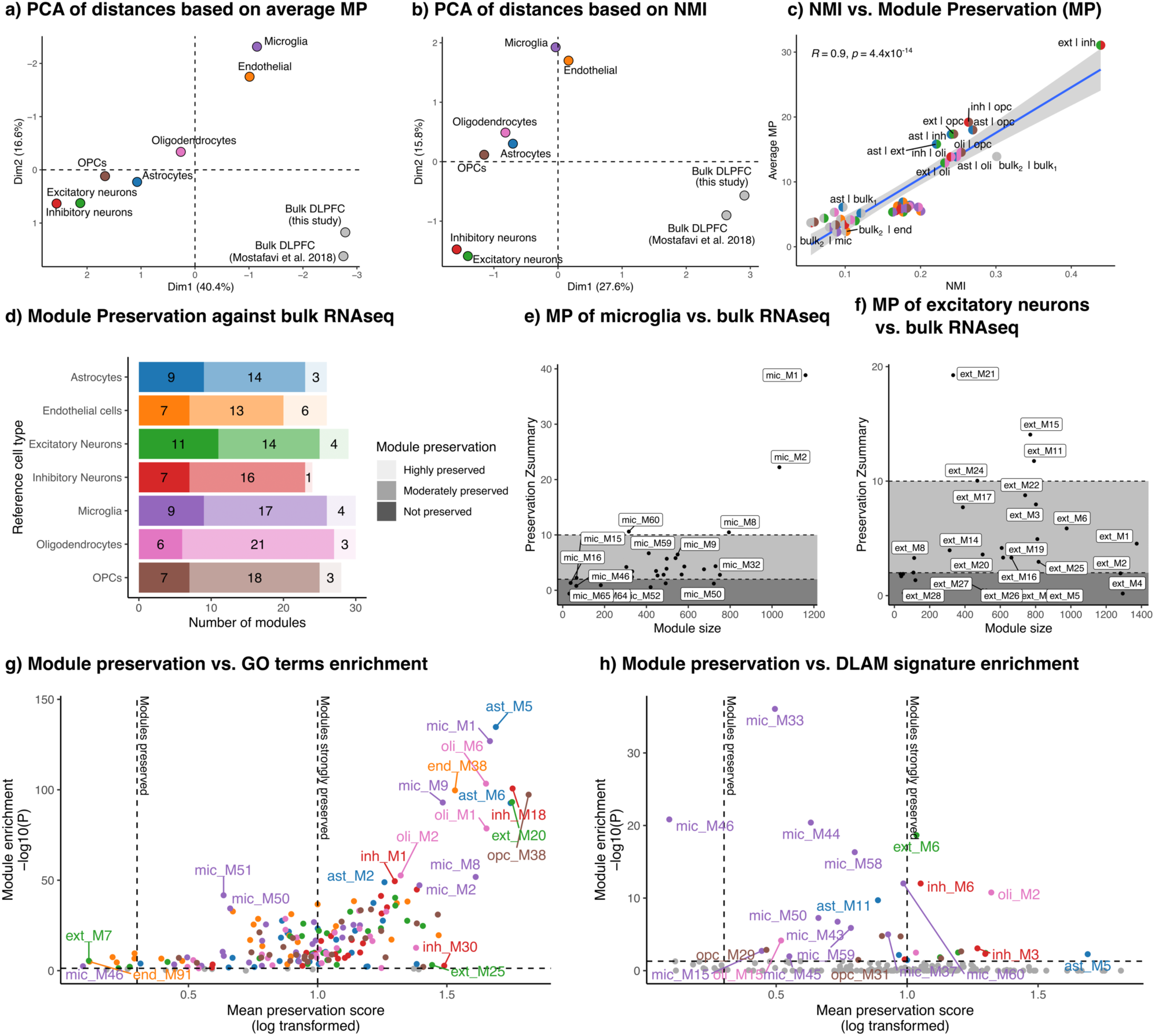
Cell type similarity modules. **a)** PCA plot showing the average module distances based on module preservation. **b)** PCA plot illustrating the module distances based on normalized mutual information. **c)** Correlation between module preservation and normalized mutual information metrics (bulk_1_ = this current study; bulk_2_ = Mostafavi et al. 2018). **d)** Module preservation results for modules from each cell type compared with bulk RNASeq. A module was considered not preserved if module preservation Zsummary < 2, moderately preserved if Zsummary ≥ 2, and highly preserved if Zsummary ≥ 10 (Langfelder et al. 2011). **e-f)** The Zsummary statistic (y-axis) as a function of module size (x-axis), for modules derived from Microglia **(e)** and Excitatory neurons **(f)**. Each dot represents one module, labeled accordingly. Dashed lines indicate the Kruskal–Wallis test cutoffs at 2 and 10. **g-h)** Comparison between average module preservation (compared with modules from other cell types and bulk RNASeq data, log_10_(average Zsummary); x-axis) and functional enrichment analysis (best -log_10_(*P*-value) for each module; y-axis). Vertical dashed lines indicate module preservation thresholds: not preserved < 2, moderately preserved if ≥ 2 and < 10, and highly preserved if Zsummary ≥ 10. Horizontal dashed lines indicate nominal *P*-value threshold of 0.05. **g)** shows results for the broad list of Gene Ontology (GO) terms, while **h)** shows results of enrichment for a selected list of damage lipid-associated microglial (DLAM) genes (Podleśny-Drabiniok et al. 2024; Gerrits et al. 2021).

To better understand the single-nucleus modules that were less preserved in comparison to other cell types and the bulk coexpression networks, we investigated their likely functional attributes. Our analysis revealed that more preserved single-nucleus modules tended to be strongly enriched for gene ontology terms, independent of cell type, as shown in **Figure 2g** (**Supplementary Tables 17-23**). The same trends were observed when comparing the preservation with bulk RNASeq alone (**Supplementary** Figure 10). However, even with a comprehensive tool like gProfiler, which integrates information from eight distinct databases, we were unable to annotate the non-preserved modules. To address this limitation, we conducted a Fisher’s exact test using a selected list of damaged lipid-associated microglia and macrophages - DLAM - genes (Podleśny-Drabiniok et al. 2024; Gerrits et al. 2021) and, surprisingly, found that some of these specific non-preserved molecular systems were significantly enriched for these genes (**Figure 2h** and **Supplementary Table 24**). These results demonstrate that the single-nucleus data, as expected, capture specific molecular systems that may have specialized cell-type-specific functions not described in public databases such as GO.

### Modules recapitulate cell subpopulation clusters/states and reveal new biological systems

To determine whether the identified coexpression modules resembled specific cell clusters or states, we compared the module genes with differentially expressed gene (DEG) markers for 95 subpopulation clusters identified in Green et al, 2024 (Green et al. 2024). We calculated Fisher’s exact test and Jaccard index to measure the gene overlaps between each module and DEG gene sets, analyzing each major cell type group separately. A Bonferroni *P*-value threshold of 0.05 was considered significant. Overall, we found many modules with significant overlaps (**Supplementary** Figures 11-13 and **Supplementary Tables 25-29**). In many cases, modules showed enrichments with multiple cell subpopulations similar to the signature DEG patterns (**Figure 3 a-f**). A few were modules enriched with specific subpopulation clusters, indicating that these modules captured specific cell communities. For example, mic_M9, a module enriched with ribosome and mitochondria genes, showed strong enrichments with microglia subpopulations involved with enhanced-redox functions such as Mic.9 and Mic.10 (adj *P*-value = 2.64x10^-76^ and 2.22x10^-72^, respectively) (**Figure 3 a, 3 d-e**). Also, the mic_M46, enriched with DLAM genes, captured the gene signatures of Mic.12 and Mic.13, two lipid-associated subpopulations (adj *P*-value = 1.51x10^-6^ and 0.005, respectively), with overlapping genes like *EYA2*, *FOXP1*, *DIRC3*, and *GPNMP* (**Figure 3 b-c**). Of note, these subpopulations express AD risk genes as *APOE, GPNMB, SPP1*, and *TREM2*.

**Fig. 3.**
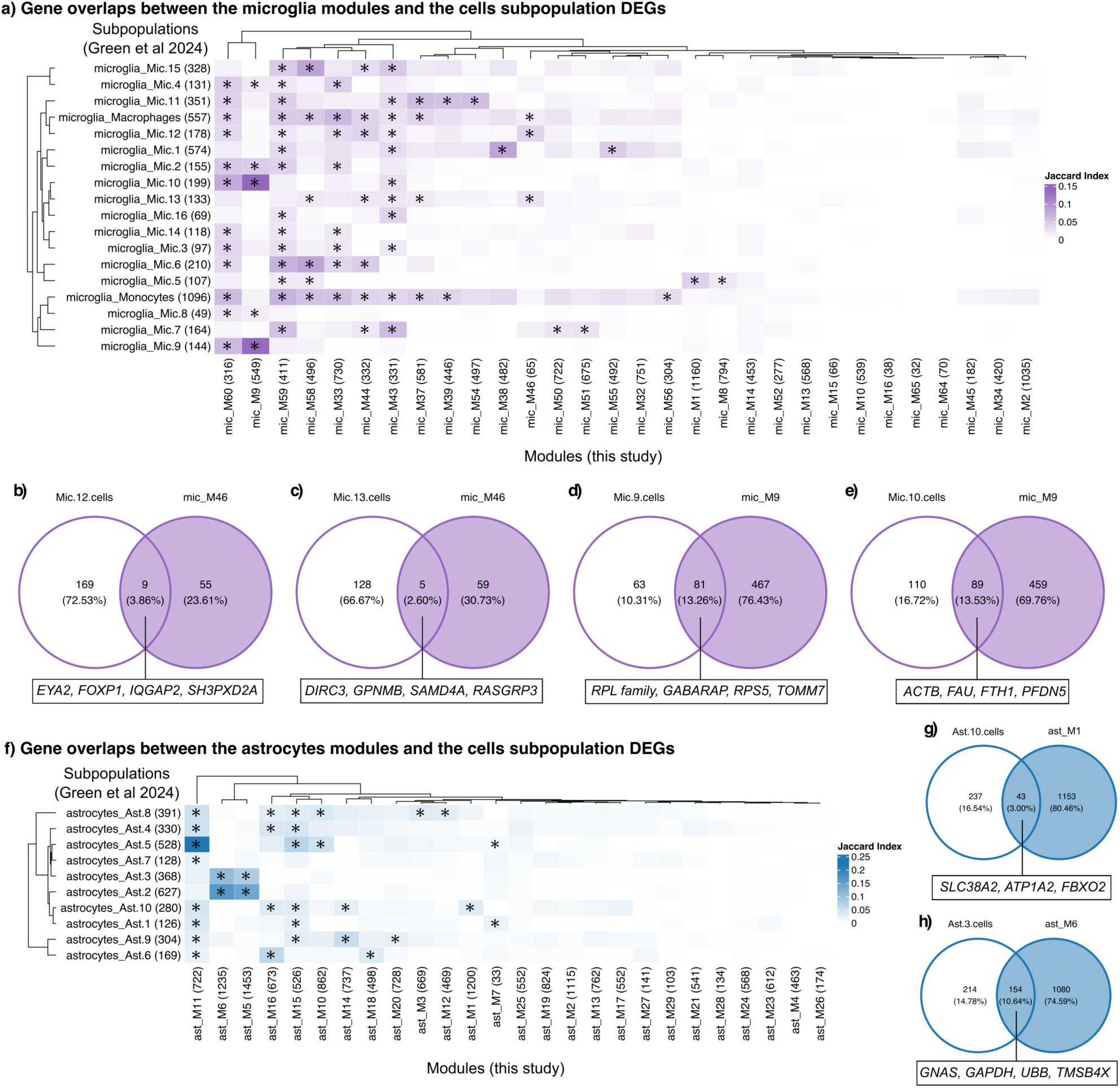
The module trait network approach captures new biological systems in the human brain. **a)** Jaccard index results for overlap analysis between the modules and the cells subpopulation types of marker genes for microglia. **b-e)** Diagrams indicating the number of genes in overlaps for selected modules and cell subpopulation type markers with significant overlap for microglia. **f)** Jaccard index results for overlap analysis between the modules and cells subpopulation types of marker genes for astrocytes. **g-h)** Diagrams indicating the number of genes in overlaps for selected modules and cell subpopulation type markers with significant overlap for astrocytes.

In the astrocyte cells (**Figure 3f**), we highlight two examples of modules also capturing signatures of cell subpopulations. The ast_M1 (enriched with genes from the Mostafavi m109, collagen biosynthesis, and chromatin organization) showed a significant overlap with marker genes of the stress response cells of Ast.10 subpopulation (adj *P*-value = 2.59x10^-5^), with overlapping genes related to synaptic transmission, such as *SLC38A2*, *ATP1A2* and *FBXO2* (**Figure 3g**). And the ast_M6 (enriched with ribosomal and mitochondrial genes) showed strong enrichments with the homeostatic cells of Ast.2 and enhanced mitophagy of Ast.3 (adj *P*-value = 1.64x10^-132^ and 7.01x10^-76^, respectively – **Figure 3h**). Although these examples show that coexpression modules can represent cell subpopulations observed from single-cell clustering approaches, not all modules captured previously annotated clusters of cells. For example, the ast_M19, enriched with nucleotide metabolism, was not enriched with makers of any annotated subpopulation clusters from astrocytes (**Figure 3f**). This result shows that our coexpression network approach captures new biological systems, not necessarily the sub-clusters found by the cell clusterization methods (i.e. Leiden) applied prior to the cell annotation.

### Identifying modules associated with ADRD phenotypes

Next, we systematically examined the associations between the modules and seven AD-related traits (**Figure 4**). These included Alzheimer’s dementia, rate of cognitive decline, cognitive resilience operationalized as cognitive decline after regressing out the effects of brain pathology, the accumulation of pathology measured separately for amyloid-β and PHFtau tangles, phosphorylated TDP-43 protein that captures LATE neuropathological changes, and a measure of global AD pathology that captures the overall burden of the disease in the aged brain (**Supplementary Table 30**).

**Fig. 4.**
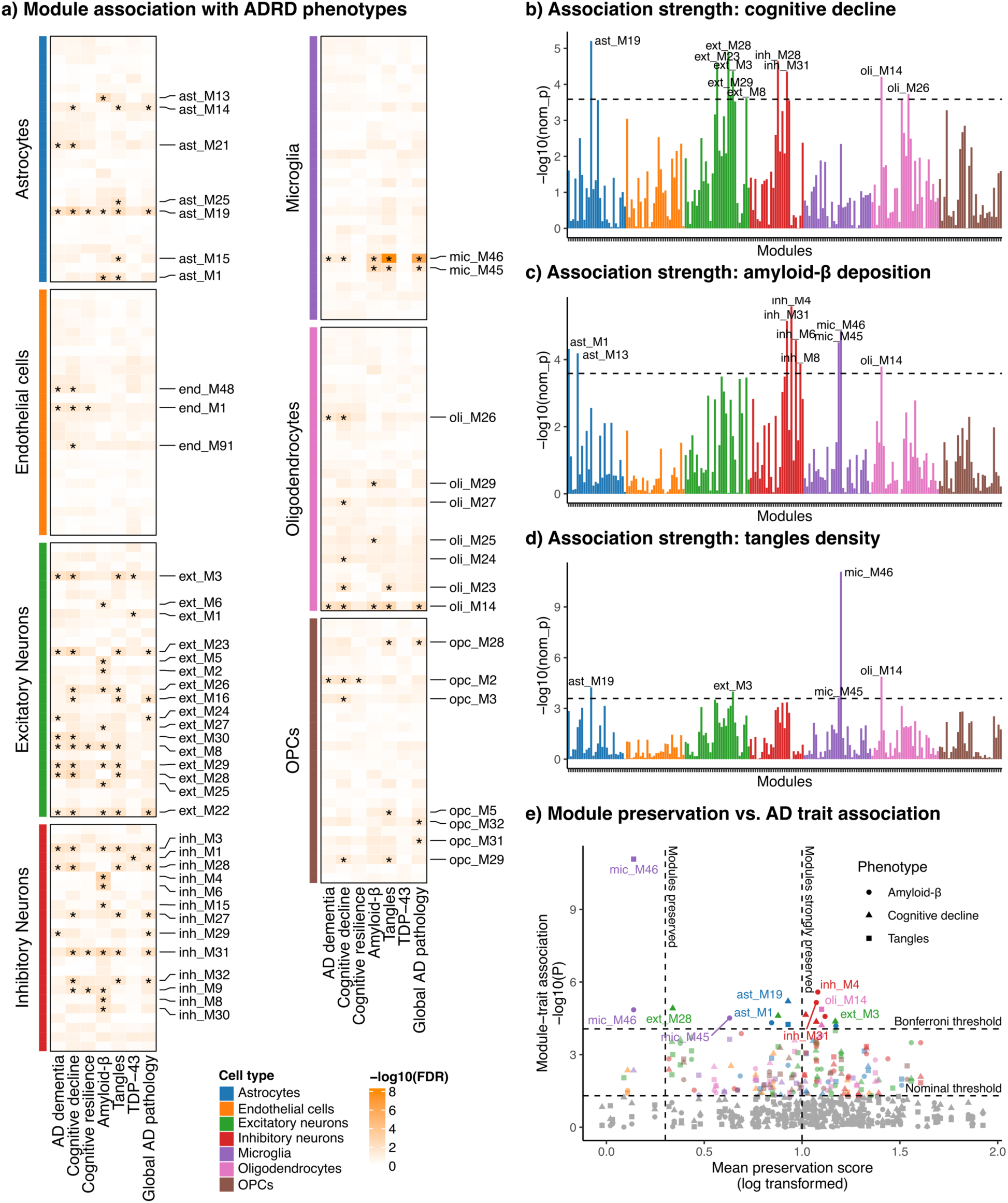
Associations between modules and AD-related traits. **a)** Heatmap showing the association results between the modules and the AD traits. The *P*-values are from linear or logistic regression models adjusted by age, sex, and years of education. Asterisks indicate that the results reached the threshold of significance (FDR adj *P*-value < 0.05). All the 193 modules are ordered according to Figure 1. **b-d)** Association strength between the modules and three AD traits: global cognitive decline **(b)**, amyloid-β deposition **(c)**, and tangle density **(d)**. The y-axis shows -log_10_(*P*-values) and dashed lines show the Bonferroni threshold (Bonferroni adj *P*-value < 0.01); only the modules that achieved the Bonferroni threshold were labeled for easy visualization. Each color is one cell type. **e)** Module preservation measures comparison with module trait association. The x-axis shows average module preservation as log_10_(average Zsummary) while the y-axis shows the best AD phenotype association as -log_10_(*P*-value) for each module. Point shapes represent different phenotypes (circle = amyloid-β; triangle = cognitive decline; square = tangles). Vertical dashed lines indicate module preservation thresholds: not preserved < 2, moderately preserved if ≥ 2 and < 10, and highly preserved if Zsummary ≥ 10. Horizontal dashed lines indicates nominal and Bonferroni adjusted thresholds (=0.05) for the module association with the three AD traits shown.

Overall, 55 of 193 modules (28%) were associated with at least one AD trait (Adj. *P-*value < 0.05) across all seven major cell types (**Supplementary** Figure 14). A higher fraction of associations was observed in modules derived from excitatory and inhibitory neurons (55% and 54%, respectively), while other cell types showed fewer modules linked to traits (**Figure 4a**). The strongest associations with global cognitive decline were found with the astrocyte module ast_M19 (*P*-value = 6.23 x 10^-6^) (**Figure 4b**). While for amyloid-β, inh_M4 showed the best association (*P*-value = 2.59 x 10^-6^) (**Figure 4c**). This module was not associated with other traits tested. The strongest association over all modules and traits was identified for the microglial module mic_M46 with PHFtau tangle density (P-value = 8.36 x 10^-12^) (**Figure 4d**). The mic_M46 was also associated with Alzheimer’ dementia, cognitive decline, and AD pathologies. As for TDP-43, only 3 modules reached significance, ext_M3, ext_M1, and inh_M1.

In our prior study of coexpression analysis in the DLPFC, we identified a particular coexpression module, m109, which showed the strongest association with a range of AD traits, including cognitive decline, AD pathology, and cognitive resilience. This finding has led to several follow-up studies focusing on specific genes within m109. Specifically, the *INPPL1* and *PLXNB1* genes were found to be related to extracellular amyloid-β in astrocyte cultures, while AK4, ITPK1, IGFBP5, and HSPB2 proteins were associated with late-life cognition, explained by microstructural changes in the brain (Mostafavi et al. 2018; Kim et al. 2019; Yu et al. 2022). Interestingly, genes involved in m109 are also co-regulated in our cell-type-specific coexpression networks. We identified five modules in oligodendrocytes (oli_M15, oli_M13, oli_M12, oli_M14 and oli_M5), four modules in astrocytes (ast_M1, ast_M13, ast_M15 and ast_M2), three in endothelial cells (end_M57, end_M58, end_M6), two in excitatory neurons (ext_M21 and ext_M16), two in inhibitory neurons (inh_M4 and inh_M19) and one in microglia (mic_M37) with significant number of overlapping genes with m109 (FDR < 0.05; Fisher exact test). As expected, some of these m109-related modules were found associated with AD traits, such as ast_M1, inh_M4 and oli_M14. However, no significant associations were observed in m109-related modules from microglia or endothelial cells, underscoring the cell-type-specific responses to disease.

Further, we then compared how the associations with AD traits related to metrics of module preservation. Importantly, as noted in the comparison of enrichments between GO terms and specialized gene sets (**Figure 2g-h**), we found the strong association of mic_M46 with PHFtau tangles contrasting with its overall low preservation compared to modules from other cell types or from bulk RNASeq (**Figure 4e**). We also found moderately preserved modules, such as ast_M19, ast_M1, and mic_M45, with significant associations with AD traits as well as modules highly preserved across cell types also displayed like inh_M4, inh_M31, and oli_M14. These findings highlight the complex interplay of molecular systems affected by AD and suggest that these less preserved modules represent critical cell-specific mechanisms to AD pathophysiology that will be missed in bulk RNASeq.

### Bayesian networks modeling the cascade effect of AD

To infer direction of effects, we modeled a Bayesian network with nodes representing either gene modules (averaged expression) or AD traits, and with arrows assigning the most likely causal direction (Tasaki et al. 2015). The manner in which these Bayesian networks reduce the number of edges versus raw correlation-based networks is important because we are looking for molecules upstream of the phenotype. Analogous to the previous work (C. Gaiteri et al. 2014), we included three levels of information: 1) Coexpressed modules highly associated with pathology and cognition. **Figure 4 b-d** shows the association results we ran for all modules, colored by cell type. From this step, 11 coexpression modules were prioritized for direct effect inference (Bonferroni *P*-value < 0.01). 2) Subpopulation of cells associated with AD. To prioritize which cells to include, we referred to the results published by Green et al. (2024), where the authors conducted a mediation analysis on subpopulations of cells and AD endophenotypes, positioning them along the AD cascade using the same snRNAseq dataset. The lipid-associated Mic.12, Mic.13, and the stress response Ast.10 cells were prioritized for the Bayesian network. 3) Phenotypes of interest. Three nodes representing AD traits were included, amyloid-β accumulation, tangles density, and global cognitive decline.

Figure 5a shows the directed acyclic graph representation for the Bayesian networks. In these networks, arrows indicate the direction of effect, edge thickness is related to the weights after 500 interactions, squares represent the modules of genes colored by their respective cell type, diamonds represent subpopulations of cells, and circles represent the AD-relevant phenotypes (**Supplementary Tables 31-32**). The edges in the network were modeled according to the direction of effect in the AD cascade as previously reported in these data, starting with amyloid-β, followed by tangles culminating in cognitive decline (David A. Bennett et al. 2004; D. A. Bennett et al. 2005; Farfel et al. 2016). In this network analysis we are looking for modules that have a direct edge to a trait. As a result, two inhibitory neuron modules, inh_M6 (enriched for protein binding and cell proliferation), and inh_M31 (enriched with protein binding and mitochondria genes) were identified upstream of amyloid-β, and at the beginning of the AD cascade, while the microglial module mic_M46, one of the modules unique to the snRNAseq coexpression networks, was directly linked to tangles density. Additionally, we found the ast_M19 to be associated with cognitive decline through the stress response cells of Ast.10, meaning that this specific population of cells is likely mediating the association of the genes in module 19 from the astrocyte networks with cognitive decline. Importantly, ast_M19 was also upstream of the modules ast_M1, inh_M4 and oli_M14, enriched for the biological signature from Mostafavi’s m109 (previously directly linked to cognitive decline). Since ast_M19 shared no genes with Mostafavi’s m109, this could possibly represent a truly new biological system captured by the snRNASeq.

**Fig. 5.**
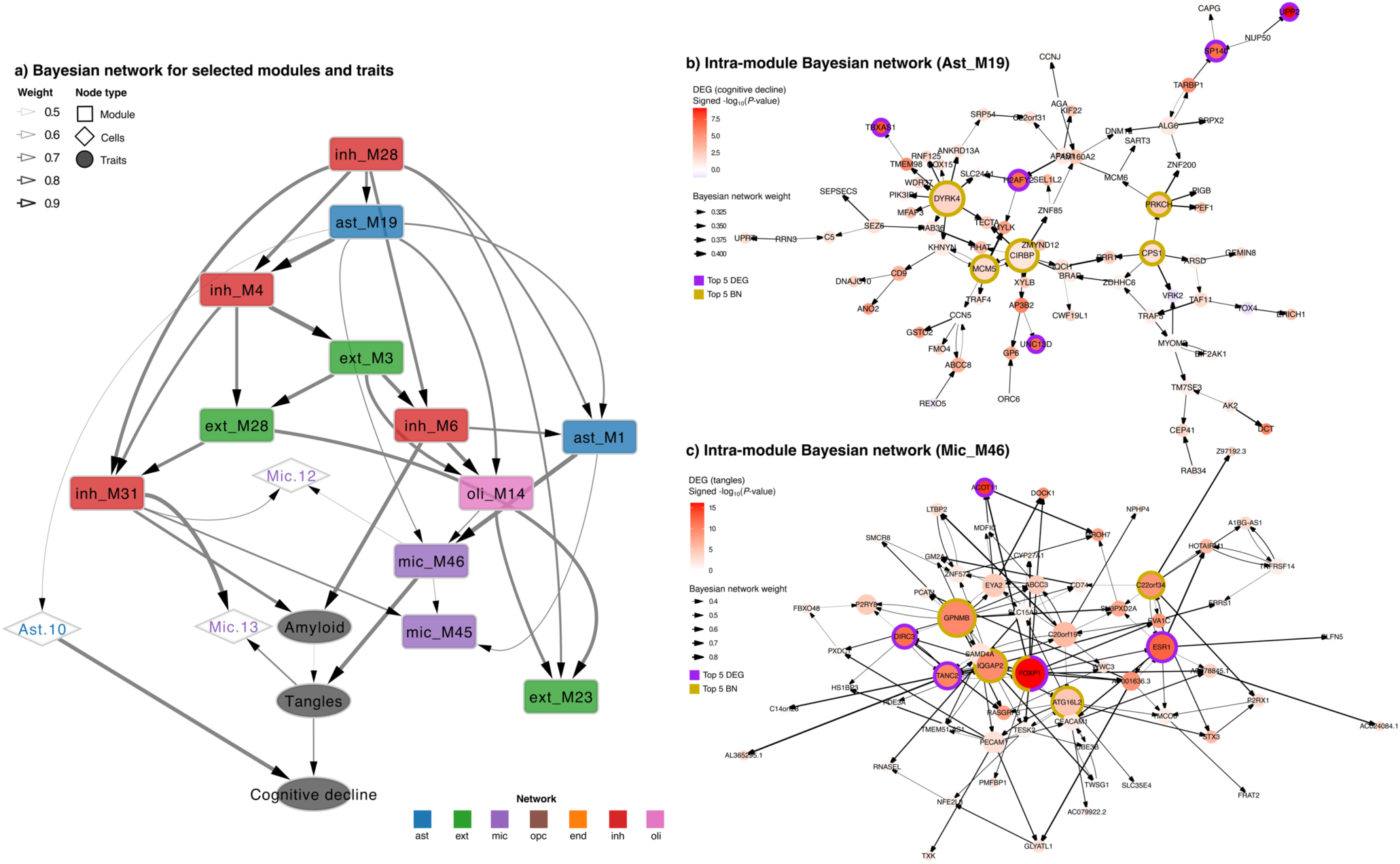
Inter- and intra-module Bayesian network analysis. **a)** A directed acyclic graph representation from the Bayesian network algorithm representing the modules, AD pathologies (amyloid-β and tangles) and cognitive decline, plus selected cell subpopulation proportions. Circles indicate AD-relevant traits, diamonds for cell subpopulation type populations, and squares for modules of genes. **b)** Top 100 genes in the ast_M19 network. Top genes were selected according to their association with AD pathology and cognitive decline measurements. **c)** All genes in the mic_M46 module. **a-c)** The arrows indicate the direction of effect, and edge thickness is related to the weights after 500 interactions. Node size is proportional to the node’s connectivity. Node internal color shades represent the strength of gene differential expression (DEG) with each respective trait, and nodes with colored strokes represent selected genes: purple = Top 5 DEG, gold = Top 5 hub genes (highest out-degree).

The module ast_M19 is composed of 824 genes/nodes. These genes were enriched for transcription factors, GO molecular functions, including ion binding, protein binding, and catalytic activity, and GO biological processes, such as metabolic and acid biosynthetic pathways. This module also contained one AD GWAS gene (the SLC2A4 regulator - *SLC2A4RG*), 2 plaque-induced ortholog genes (cathepsin L - *CTSL* and a member of the transmembrane family - *CD9*) (Chen et al. 2020), 58 genes from the mitochondria GO list (e.g. *COX15, CASP9, NUDT2*) and 43 genes from GO lipid metabolism (e.g. *DDHD2, NRG4, PIK3IP1*). To determine how much of its expression was explained by astrocyte subpopulations, we ran a linear regression with the module’s average expression and the astrocyte cell fractions. We found that the ast_M19 had its expression partially explained by the stress response Ast.10 (34%); the enhanced mitophagy Ast.3 (9%); the homeostatic-like Ast.1 (7%); and to a less extent by the Ast.7, Ast.8 and Ast.9 cells (**Supplementary** Figure 15). When we unpack the modules with a direct edge to a trait, we are most interested in network drivers which tend to be the most connected node in the module. To explore the ast_M19, we modeled an intra-network between genes analysis of ast_M19. (**Supplementary Table 33**). Figure 5b shows the top 100 genes (ranked by their association with AD traits) of this specific module. The dual specificity tyrosine phosphorylation regulated kinase 1A (*DYRK4*) gene seemed to be a master regulator of at least 10 other genes (e.g. *MFAP3, TMEM98, COX15*). Cold-inducible mRNA binding protein (*CIRBP*) gene was directly regulating other 5 neighbors (e.g. *ZNF85, XYLB, TECTA*), and the carbamoyl-phosphate synthase 1 (*CPSI*) gene was directly linked to *PRKCH, PRR11, ARSD, VRK2* and *ZDHHC6*.

The module inh_M6 was also upstream of m109 related systems (ast_m1 and oli_m14) and was directly linked to amyloid-β. The inh_M6 is a neuronal module with 706 genes enriched with several molecular functions and biological processes, including protein binding, kinase and transferase activity, cellular and phosphorus metabolic processes regulation, neurogenesis, and neuron differentiation (**Supplementary Table 34**). This module also had 4 AD GWAS genes (*PLCG2, SNX1, NCK2* and *JAZF1*), 79 genes from the cell proliferation GO list (e.g. *TMEM131L, STAT5B, IL20RB*), 52 genes from proteolysis (e.g. *DLC1, IFT52, USP4*), and 13 from lysosome (e.g. *ATP11A, PLBD2, SNX1*). **Supplementary** Figure 16 shows the acyclic graphic for the network built for the top 100 genes associated with AD traits within inh_M6. The analyses revealed the following genes as master regulators of their surroundings: *ATP11A, GAB2, USP28,* and *RBM5*.

In order to make the modules identified in the Bayesian networks experimentally tractable, we generated intra-module Bayesian networks for the mic_M46 (Figure 5c and **Supplementary Table 35**). This module was downstream of the modules ast_M19, ast_M1 and oli_M14 (last two Mostafavi’s m109 enriched) and directly associated with tangles accumulation. The mic_M46 is a module composed of 65 genes and enriched with DLAM signature genes. The network revealed the *FOXP1* upstream of other 6 genes, including *TESK2, DIRC3* and the long-non-coding *AC078845.1*. The PD risk gene *GPNMB* (up in disease associated microglia - DAM cells (Keren-Shaul et al. 2017)) was regulating *SAMD4, CD74, ABCC3, TMEM51* and *P2RY8*.

## Discussion

Here, we investigated the aging human brain networks by leveraging bulk and snRNASeq data from ROSMAP participants. By applying the module-trait network approach, we built coexpression modules for the seven major cell types of the DLPFC. These modules were deeply characterized, from functional annotation to preservation analysis. We found modules specific to single nucleus datasets that were not present in bulk RNASeq datasets and uncovered cell-specific biological systems and genes relevant to AD traits. We also showed that our approach captured modules that potentially represent cell subpopulations but also captured new biological systems. Finally, by integrating three layers of information—phenotype data, modules of genes, and cell subpopulations—we performed inference models to prioritize modules directly linked to AD phenotypes and important gene regulators in these networks.

Previous studies on gene coexpression modules using bulk brain tissues were pivotal to understanding molecular mechanisms and functions related to AD by revealing important genes and modules. For example, the discovery of *TYROBP* as a key regulator in a microglia-specific module associated with AD (Zhang et al. 2013); and the prioritization of module 109, linked to cognitive decline (Mostafavi et al. 2018) and the following assessment of the role of AK4 at the protein level (Yu et al. 2022, 2018). At the cell-specific level, however, studies of coexpression networks have been historically limited to small sample sizes, with most findings focused on identifying cell-specific coexpression signatures rather than finding their relationship with AD. Also, there are still debates on methods for inferring gene coexpression from single-cell data. With methods designed to model coexpression from single-cell sparse data (Su et al. 2023; Lopez-Delisle and Delisle 2022; Wang, Choi, and Roeder 2021; Mathys et al. 2024) to others methods, such as hdWGCNA (Morabito et al. 2023) and our approach, that are based on aggregation strategies, designed to create pseudo bulk matrices and apply traditional methods originally designed for bulk RNASeq data. Similarly to our findings, most previous studies observed a large proportion of module conservation between cell types, suggesting consistent regulatory landscapes across cells (McKenzie et al. 2018; Harris et al. 2021; Morabito et al. 2023). More recently, a well-powered study, also derived from ROSMAP samples (Mathys et al. 2024), was able to identify and annotate at a finer degree modules representing specific populations of cells. Their results also showed cell-specific modules related to AD pathology, including enrichments of modules with pathology-associated astrocytes and gliosis, modules shared in microglia and astrocyte cells. Similarly to our findings, these results can help improve our understanding of gene regulation in the cell and disease-specific context. Our study, however, advances the field by incorporating specialized cell subpopulations into direct effect inference, providing novel insights into the molecular systems underlying AD.

In our analysis, by discovering coexpression modules separately for each data modality, we were able to measure if the clusters of genes were preserved across the networks. As expected, we observed that modules tend to be more similar between groups of related cells, such as between excitatory and inhibitory neurons or microglia and endothelial cells. And that, overall, modules derived from snRNASeq are quite distinct from modules derived from bulk RNASeq (Figures 2a-f). We also found that modules highly preserved tended to be more enriched for common molecular functions and biological processes as described in the Gene Ontology (Figures 2g-h). However, in the cases where modules were less or not preserved in other cell types, their functions are more likely to be more specific to a certain cell population or condition. This was the case noted for the module mic_M46. This module was not particularly enriched for GO terms, but instead, it was enriched for microglial genes linked with response to lipid and amyloid-β in AD context (named DLAM) (Gerrits et al. 2021; Podleśny-Drabiniok et al. 2024). In addition, we showed that our modules not only can capture signatures of annotated cell populations (e.g., mic_M46 and the subpopulations annotated as Mic.12 and Mic.13) but also can discover new biological systems (e.g., ast_M19) not previously identified by the single-cell clustering (Figures 3 and **Supplementary** Figures 10-12). Providing additional insights into cell-specific gene-gene relationships.

Our association results highlighted several biological systems related to AD traits. Among the strongest associations, we identified the module ast_M19 associated with decline in cognition, inh_M4 associated with amyloid-β, and mic_M46 associated with pathology accumulation (Figure 4). Importantly, mic_M46 was not preserved in bulk RNASeq data, while the others were only moderately preserved, supporting the novelty of our findings from deriving coexpression networks from snRNASeq. Finally, by modeling the direction of relationships between modules, traits, and cell populations via Bayesian network approach, the inh_M6 was upstream of amyloid-β, the mic_M46 directly associated with tangles accumulation, and the ast_M19 associated with cognitive decline through the mediation of the Ast.10 subpopulation of cells (Figure 5). Furthermore, we also modeled the intra-module direct effect to highlight key genes in these modules, such as *DYRK4* in ast_M19 and *FOXP1* in mic_M46, offering a foundation for targeted experimental validation and therapeutic exploration.

The study has many strengths. First, ROSMAP participants are deeply phenotyped over years prior to death and follow-up participation and autopsy rates are very high reducing bias. Further, they are recruited without dementia in the community eliminating referral bias that typify clinic-based studies (Schneider, Aggarwal, et al. 2009). All pathologic data is collected blinded to clinical data. Third, the study is sufficiently large that we can find small, unique, but stable cellular communities. Our work also has limitations. The snRNASeq experiment captures the RNA from the nucleus, not capturing the cytoplasm RNA gene expression (Thrupp et al. 2020). Thus, we also compared the snRNASeq results with constructed coexpression networks from bulk RNASeq, where the experiment captures RNA from the complete cell. Regarding the functional annotations, it is also important to highlight that many of the modules associated with AD traits are multi-functional. Not often, these biological systems are enriched with a variety of gene ontology terms, can be shared across distinct networks, and play distinct roles depending on the context where they are placed. From the functional enrichment analysis, we aimed to prioritize a single functional term for practical purposes (**Supplementary Tables 17-23**). However, we acknowledge that this is a non-trivial task, and the complete summary statistics (available on GitHub) should be considered for post-hoc analysis. In addition, the ROSMAP participants are old, non-Latino White individuals with high levels of education and, therefore, not representative of the general population. Also, although the sample size is adequate for the methodologies presented here, by increasing the number of participants with more diverse genetic ancestry backgrounds, we hope to be able to discover new subpopulations of cells that may drive AD trait associations.

To our knowledge, this is the first study to include specialized cell subpopulations for direction of effects in a Bayesian framework. Our findings showed that these cells potentially mediate the interaction of a cluster of genes with a disease phenotype and may provide insights and new directions for targeted experiments highlighting once again the importance of studying molecular systems and deriving gene modules in a cell-specific and pathology/disease condition context.

## Methods

### Clinical evaluations

The participants were from two prospective studies of aging and dementia, the Religious Orders Study (ROS) or the Rush Memory and Aging Project (MAP), commonly referred to as ROSMAP. Both are cohort studies of risk factors for dementia, AD, and other aging outcomes. ROS enrolls older catholic priests, nuns, and monks across the USA while MAP enrolls older laypersons from the greater Chicago metropolitan area, Illinois (David A. Bennett et al. 2018). The participants agreed to annual detailed clinical evaluations and organ donation at the time of death. They also enrolled without known dementia, provided written informed consent, and signed an Anatomical Gift Act. A Rush University Medical Center Institutional Review Board approved each study. Clinical evaluations are administered annually by testers blinded to data from previous years. The cognitive battery examination contains 21 tests, 19 of which were used to construct a global composite measure of cognitive decline used in this study. Raw scores for individual tests were standardized using the baseline means and standard deviations of the entire cohorts, and then averaged across the tests to obtain the composite score (Boyle et al. 2018; Robert S. Wilson et al. 2015). To assess person-specific cognitive resilience, the rate of change in global cognition over time was estimated, controlling for demographic and pathology variables (De Jager et al. 2012).

### Neuropathologic evaluations

At autopsy, the brain was removed, weighed, and sectioned coronally into 1 cm slabs. One hemisphere was frozen at -80°C for biochemical studies, while the other was fixed in 4% paraformaldehyde for neuropathologic evaluations (David A. Bennett et al. 2005; Schneider, Arvanitakis, et al. 2009). The average post-mortem interval was 8.4 hours (SD=6.0). The evaluations were conducted by examiners blinded to all clinical data, ensuring an unbiased examination. The post-mortem assessments included neuropathologies of AD and limbic-predominant age-related TDP-43 encephalopathy (LATE) with a standard protocol (Boyle et al. 2021). The pathological diagnosis of AD was based on the National Institute on Aging Reagan criteria. Bielschowsky silver stain was used to visualize neuritic plaques, diffuse plaques, and neurofibrillary tangles, including five areas of the brain: frontal, temporal, parietal, entorhinal, and hippocampal cortices (Schneider, Arvanitakis, et al. 2009). The 15 counts obtained in this step were scaled and used to measure the global AD pathology burden, for each individual (David A. Bennett et al. 2006). Then, unbiased methods were employed to quantify the accumulation of Aβ plaques and paired helical filaments (PHF) tau tangles, enabling access to specific indices of AD pathology (R. S. Wilson et al. 2007). Briefly, images were captured from the outlined tissues using Stereo Investigator software version 9 and an Olympus BX-51 microscope equipped with a motorized stage. A grid was placed over the area, and approximately 25– 50% of the brain region was sampled. After camera and illumination calibration, images at each sampling point were acquired with the motorized stage. Aβ quantification was performed via image processing, based on approximately 90 images from the cortex for larger brain regions and about 20 images from smaller regions (e.g., the hippocampus). The mean Aβ fraction per brain region and per subject was calculated. Details were previously described (Kapasi et al. 2021; R. S. Wilson et al. 2007).

The LATE neuropathology indices were assessed using immunohistochemistry. TDP-43 staining was performed on six brain regions: the amygdala, hippocampus, dentate gyrus, entorhinal cortex, midfrontal cortex, and middle temporal cortex. Semiquantitative measures of pathogenic TDP-43 were analyzed in neurons and glia within a 0.25 mm² area of highest density. Finally, four stages were defined based on the pathological distribution of TDP-43: 0 = no TDP-43 lesions, 1 = TDP-43 localized to the amygdala, 2 = TDP-43 extending to the hippocampus or entorhinal cortex, and 3 = TDP-43 extending into the neocortex (Kapasi et al. 2020).

### snRNAseq data processing and QC

Previous publications have extensively described the ROSMAP snRNASeq and this is a brief description of data processing (Fujita et al. 2024; Green et al. 2024). Nuclei were isolated from 479 DLPFC human brain tissues. The tissues were processed as 60 batches, and each batch consisted from 8 participants. In each batch, nuclei suspension of 8 participants were mixed together, and single-nucleus RNASeq library was prepared using the 10x Genomics 3 Gene Expression kit (v3 chemistry). The libraries were sequenced, and read mapped and UMI counting were performed using CellRanger v6.0.0 (Zheng et al. 2017) with GENCODE v32 and GRCh38.p13. Original participants of droplets in each batch were inferred by comparing SNPs in RNA reads with ROSMAP whole genome sequencing (WGS) VCF files using genetic demultiplexing software demuxlet (Kang et al. 2018). As a quality control, genotype concordance of RNA and WGS, sex check, duplicated participants, WGS QC, and sequencing depth were assessed, and 424 participants passed the QC. To annotate cell types based on single nucleus expression, nuclei were classified into 7 major cell types, and each major cell type group was analyzed separately. A total of 96 subpopulations of cells were annotated based on marker genes (syn52363764; (Fujita et al. 2024; Green et al. 2024)). Doublets were removed using DoubletFinder, and cells were clustered using Seurat (Stuart et al. 2019).

For coexpression network inference, we created pseudo-bulk matrices by summing counts per participant. The genes were filtered separated by cell type, keeping the ones with at least 1 count per million (CPM) in 80% of samples. Different methods of normalization were tested, and TMM was applied with voom for the final matrices (Law et al. 2014). For the comparison between modules and subpopulation marker genes (Figure 3), the lists of DEGs were derived from slightly different annotations using the same dataset (syn53694215; (Fujita et al. 2024; Green et al. 2024)).

### Bulk RNASeq data quality control analysis

RNA was extracted from the DLPFC using the Chemagic RNA tissue kit (Perkin Elmer, CMG-1212). The RNA quality number (RQN) was calculated using FragmentAnalyzer (Agilent, DNF-471) and it is concentration determined using Qubit broad range RNA assay (Invitrogen, Q10211) according to the manufacturer’s instructions. A total of 500 ng total RNA was used for RNASeq library generation and rRNA was depleted with RiboGold (Illumina, 20020599). A Zephyr G3 NGS workstation (Perkin Elmer) was utilized to generate TruSeq stranded sequencing libraries (Illumina, 20020599) with custom unique dual indexes according to the manufacturer’s instructions. The libraries were normalized for molarity and sequenced on an Illumina NovaSeq 6000 at 40 to 50 million reads, 150 bp paired-end. With the mRNA sequenced, we used three parallel pipelines, an RNASeq quality control (QC) pipeline, a gene/transcripts quantification pipeline, and a 3ʹ-UTR quantification pipeline. First, the paired-end sequences were aligned by STAR v2.6 (Dobin et al. 2013) to a human reference genome and annotated with GENCODE (Release 27 GRCh38). The metrics from Picard tools were analyzed for QC, and for quantification, the transcript raw counts were calculated using Kallisto (v0.46) (Bray et al. 2016). 17,294 genes were expressed in > 50% of the samples and had at least 10 counts. For normalization, conditional quantile normalization (CQN) was applied to adjust for guanine-cytosine (GC) content and gene length. The gene count matrix was converted to log_2_(CPM) followed by quantile normalization using the voom lima function (Law et al. 2014). Finally, the matrices were adjusted to remove technical confounders and the linear regression model included variables of post-mortem interval, sequencing batch, RQN, total spliced reads reported by STAR aligner, and quality metrics reported by Picard and Kallisto.

### Coexpression networks

To find the coexpression networks of the brain, we used the Speakeasy algorithm version 1.0 Matlab based (Chris Gaiteri et al. 2015). The Speakeasy uses a dynamic label propagation approach to robustly identify clusters in biological or at any given dataset, as proven by achieving the quality required by the Lancichinetti-Fortunato-Raddichi benchmarks. Another advantage of the Speakeasy is the lack of parametrization, meaning that the modules were found in an unbiased way without tuning settings for “reasonable or desirable” results. This method was also used before with ROSMAP RNASeq data and extensively compared with the highly cited Weighted Gene Co-expression Network Analysis (WGCNA) tool (Mostafavi et al. 2018).

As input for the coexpression networks, we used the gene expression matrices already normalized and adjusted. After the consensus clustering results from 100 initializations, we found the following number of modules with at least 30 genes by each cell type: astrocytes (26), microglia (30), excitatory neurons (29), inhibitory neurons (24), oligodendrocytes (30), endothelial cells (26), OPCs (28), and bulk DLPFC (34). Microglial and endothelial had 95% of their expressed genes clusterized (out of 14,395 and 8,412 genes, respectively), and all the other coexpression networks had 99% (out of > 15,000) of their genes assigned to a module. The module assignments are in the **Supplementary Tables 4-10**. Module average expression and eigengene, defined as the first principal component of the module expression, were computed for module summarization.

### Association analysis description

We ran linear and logistic regressions using the R packages lme4 (Bates et al. 2015) and lmerTest (Kuznetsova, Brockhoff, and Christensen 2017) to test the associations between the modules and the AD traits. The module average expression was used as the predictor, the phenotype matrix with the AD traits was defined as the outcome, and the following covariates were used for adjustment in the models: age at death, sex, and years of education. The *P*-values were adjusted by Bonferroni and considered significant if adj *P*-value < 0.05. We calculated the -log_10_(*P*-value) of the association for the heatmap in Figure 4.

### Gene-set enrichment analysis

Each module was carefully annotated using three approaches: **First, using the gprofiler tool** (Kolberg et al. 2023). This tool maps sets of genes to known functional information from 8 distinct datasets, including Gene Ontology (GO) database; Kegg, Reactome, and Wikipathways; regulatory motif matches from TRANSFAC; tissue specificity from Human Protein Atlas (HPA); protein complexes from CORUM, and human disease phenotypes from Human Phenotype Ontology (WP). The gost R function was used with the following parameters: organism = “hsapiens”, correction_method = “gSCS”, significant = TRUE, user_threshold = 0.05. The full results are available online on the GitHub page. **Second, using Fisher exact tests with specific gene lists.** Sometimes, the functions assigned are not informative due to poor database annotation or lack of experimental functional characterization. To overcome this issue, we downloaded lists of genes relevant to the AD phenotype, including GO-curated lists (e.g., genes of the lipid metabolism, phagocytosis, response to cytokines) (Ashburner et al. 2000; Gene Ontology Consortium et al. 2023), lncRNAs from the Gencode v37, GWAS AD risk genes (Bellenguez et al. 2022) and, plaque-induced genes (Chen et al. 2020). The Rummagene online interface was used to scan gene lists from supplementary materials (Clarke et al. 2024). The results were adjusted by Bonferroni and considered significant if adj *P*-value < 0.05. The results are available online on the GitHub page. Finally, **in the third approach, we manually checked each module.** To facilitate module interpretation, “one term” representative of the module’s function was given based on the first and second approaches described here (**Supplementary Tables 17-23**).

### Accessing module preservation across the networks

We performed a module preservation analysis to assess the shared (or distinct) modules across nine coexpression networks (Langfelder et al. 2011). The comparisons were pairwise for all seven single-nucleus networks, plus two from bulk RNASeq data: one previously published (n=478) (Mostafavi et al. 2018) and our updated version with a larger sample size (n=1,210). First, we used a permutation test to randomly permute the gene-module assignments in the test data. From this step, we estimated the mean and variance of the preservation statistic under the null hypothesis of no relationship between the module assignments in reference and test data. Fifteen distinct preservation statistics were observed (e.g., module density, mean expression, connectivity), and each was standardized considering the mean and variance, defining a Z statistic for each preservation measure. With these assumptions, each Z statistic follows the normal distribution if the module is not preserved. For comparison across the networks, we accessed the composite preservation measure defined by the following equation:

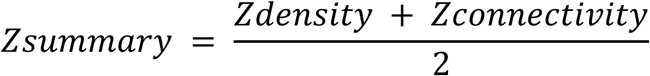

If Zsummary < 2, the module is considered not preserved; between 2 and 10, the module is considered preserved, and > 10 is highly preserved.

For the normalized mutual information, we loaded the R library “aricode” with default parameters.

### Bayesian network inference

A Bayesian network is a probabilistic graphical model that represents a set of random variables and their conditional dependencies using a directed acyclic graph. Here, we applied an ensemble Markov Chain Monte Carlo (MCMC) approach (Tasaki et al. 2015) to estimate the directed acyclic graph that describes the relationship between three types of variables: modules, cell type proportions, and AD traits. More specifically, we included: 1) the average expression of the 11 coexpression modules associated with amyloid-β, tau-tangles, and cognitive decline (at Bonferroni *P*-value < 0.01); 2) fractions of three subpopulations of cells previously associated with AD endophenotypes, Mic.12, Mic.13, and Ast.10 (Green et al. 2024); and 3) ROSMAP phenotype data for amyloid-β, tangles accumulation, and person-specific rate of decline in cognition.

A total of 500 independent MCMC chains were run using the “M.REV50” algorithm, each initialized with a different starting network structure. The edge weights represent the proportion of MCMC chains in which the edges appeared. Edges from tangles to amyloid were not allowed in the model to follow the AD cascade as previously reported in these data, starting with amyloid-β, followed by tangles, and finally, cognitive decline (David A. Bennett et al. 2004; D. A. Bennett et al. 2005; Farfel et al. 2016). A cut of edges with weight > 0.4 was applied for visualization.

From the inter-module Bayesian network results we prioritized the constructions of intra-module Bayesian for the modules ast_M19, inh_M6 and mic_M46. The module ast_M19 has 824 and the inh_M6 has 707 genes, so we selected the top 100 associated with amyloid-β, tangles accumulation, and cognitive decline; since mic_M46 had only 65 genes, we included all in the analysis. The intra-module networks were inferred with 1,000 independent MCMC chains with “M.REV50” algorithm. A cut of weight > 0.3 was set for visualization. Since the directed acyclic graph enables us to rank genes based on their number of conditional dependent connections to other genes in the module, we highlighted genes with the highest number of downstream connections as well as genes most associated with the respective traits.

## Data availability

The expression data used in this manuscript is available with the following accession numbers: ROSMAP bulk DLPFC (syn3388564) and snRNASeq (syn31512863). The phenotype data can be requested at the RADC Resource Sharing Hub at www.radc.rush.edu or the AD Knowledge Portal (https://adknowledgeportal.org). The supplementary materials include all the network analyses, including functional enrichment analysis, module prioritization, and differential gene expression analysis.

## Code availability

The relevant code generated by this study is available on the GitHub repository https://github.com/RushAlz/sn_networks_ROSMAP.

## Acknowledgements

We thank all the ROSMAP participants and the investigators and staff at the Rush Alzheimer’s Disease Center. This work has been supported the following National Institute on Aging (NIA) grants: P30AG10161, P30AG72975, R01AG15819, R01AG17917, U01AG46152, U01AG61356 and U01AG079847. This work was done as part of the National Institute of Aging’s Accelerating Medicines Partnership for AD (AMP-AD).

## Author contributions

D.A.B and S.T conceived and supervised the study. G.G, M.F, V.M, P.L.J and N.H provided the snRNASeq atlas. Y.W sequenced the bulk RNASeq samples. K.P.L applied the MTN approach across all the networks with input from C.G and S.T. The BN was modeled by R.V with input from S.T. K.P.L, S.T and D.A.B wrote the manuscript with feedback from all co-authors. All authors read and approved the manuscript. D.A.B., P.L.J., J.A.S., and C.G. funded the project.

