## Supplementary figures for "Gene module-trait network analysis uncovers cell type specific systems and genes relevant to Alzheimer’s Disease"

### List of Supplementary Tables

**Table 1:** Phenotype data for all the individuals used in the study (n = 1,210 unique donors).
**Table 2:** Description of study participants with bulk RNASeq (n = 1,210).
**Table 3:** Description of study participants with snRNASeq (n = 424).
**Table 4:** Module assignment for the astrocytes co-expression networks.
**Table 5:** Module assignment for the endothelial co-expression networks.
**Table 6:** Module assignment for the excitatory neurons co-expression networks.
**Table 7:** Module assignment for the inhibitory neurons co-expression networks.
**Table 8:** Module assignment for the microglial co-expression networks.
**Table 9:** Module assignment for the oligodendrocytes co-expression networks.
**Table 10:** Module assignment for the OPCs co-expression networks.
**Table 11:** Module assignment for the dorsolateral pre frontal cortex (DLPFC) co-expression networks.
**Table 12:** Summarized functional enrichment analysis (FEA) for module characterization. Network = DLPFC.
**Table 13:** Full summary statistics of the regression analysis between the DLPFC modules and the AD traits.
**Table 14:** Normalized mutual information matrix results.
**Table 15:** Full summary statistics of module preservation (MP) analysis across the 9 networks tested.
**Table 16:** Zsummary statistcs from the MP analysis between the single-nucleus modules and the bulk DLPFC (n=1,210).
**Table 17:** Summarized Functional Enrichment Analysis (FEA) for module characterization. Network = astrocytes.
**Table 18:** Summarized FEA for module characterization. Network = microglial.
**Table 19:** Summarized FEA for module characterization. Network = excitatory neurons.
**Table 20:** Summarized FEA for module characterization. Network = inhibitory neurons.
**Table 21:** Summarized FEA for module characterization. Network = oligodendrocytes.
**Table 22:** Summarized FEA for module characterization. Network = OPCs.
**Table 23:** Summarized FEA for module characterization. Network = endothelial cells.
**Table 24:** Fisher exact test between the modules and the damaged lipid-associated microglia and macrophages (DLAM).
**Table 25:** Jaccard similarity results for the astrocyte modules versus astrocyte cells subpopulations.
**Table 26:** Jaccard similarity results for the microglia modules versus microglial cells subpopulations.
**Table 27:** Jaccard similarity results for the excitatory neuron modules versus excitatory neurons subpopulations.
**Table 28:** Jaccard similarity results for the inhibitory neuron modules versus inhibitory neurons subpopulations.
**Table 29:** Jaccard similarity results for the oligodendrocytes modules versus oligodendrocytes subpopulations.
**Table 30:** Full summary statistics of the regression analysis between the module and the AD traits.
**Table 31:** Full results for the single-nucleus Bayesian network (BN).
**Table 32:** Filtered results for the single-nucleus BN. We keep edges with minimum 3.3 of weight, after 500 permutations.
**Table 33:** Full summary statistics for the intra-BN of ast_M19.
**Table 34:** Full summary statistics for the intra-BN of inh_M6.
**Table 35:** Full summary statistics for the intra-BN of mic_M46.

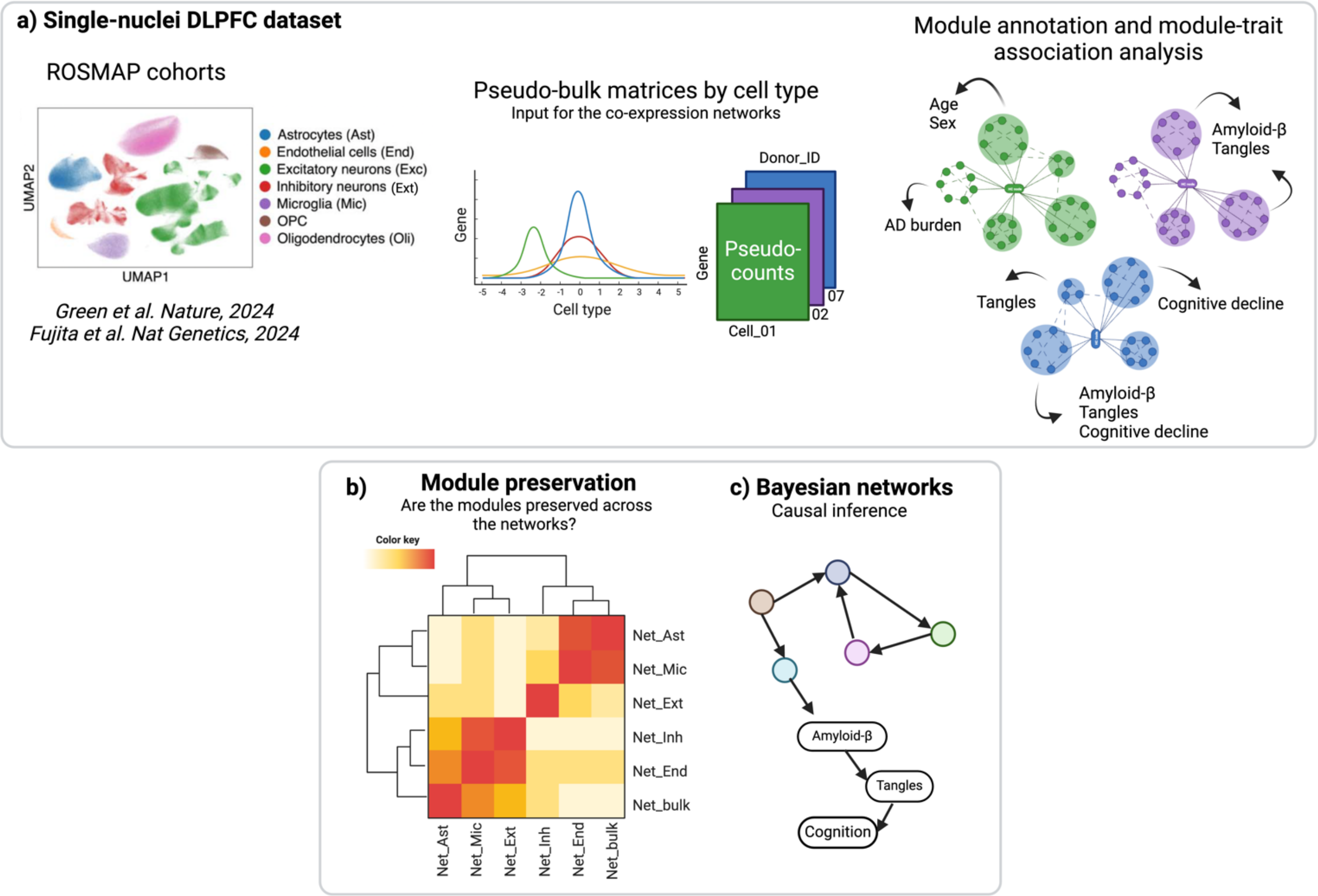

**Supplementary Figure 1: A graphic overview of the module-trait network (MTN) approach.**
a) The ROSMAP snRNASeq data used for the networks covers 1.6 million cells, 7 major cell-type groups and 96 sub-cell types (Fujita et al. 2024; Green et al. 2024). Pseudo-bulk matrices were created and used as input for the coexpression networks. The modules were annotated according to their biological functions and their average expression used for trait-association analysis. b) Pairwise measures of module preservation were calculated to identify specific or shared systems across all the networks. c) Bayesian networks were modeled for causal inference (Tasaki et al. 2015). Figure generated with biorender.

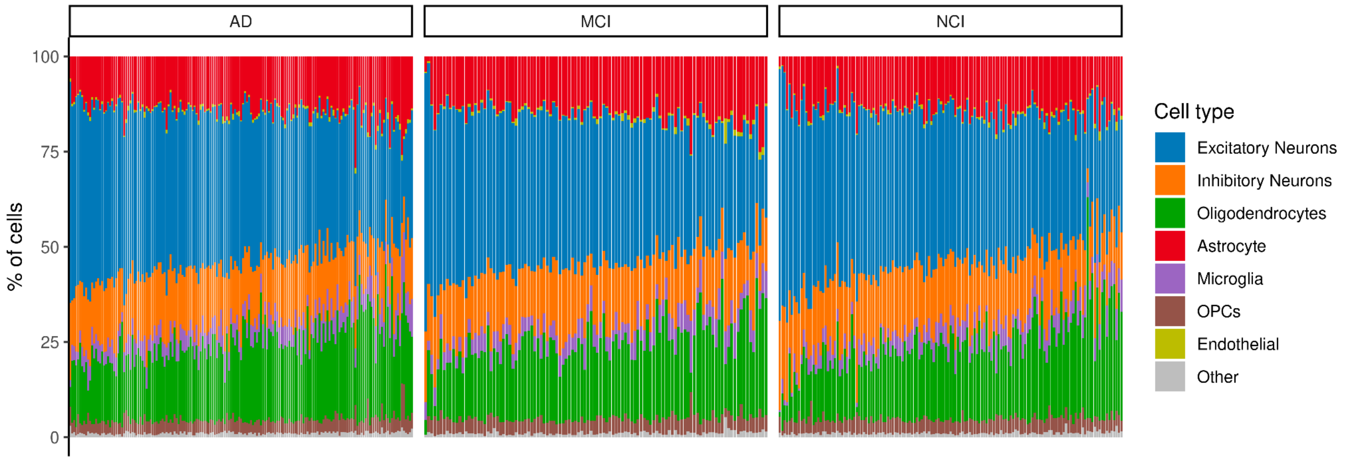

**Supplementary Figure 2: The percentage of single-nucleus cells by individual.**
Excitatory neurons accounts for 39.54% of the DLPFC cells, followed by Inhibitory neurons: 15.68%, Oligodendrocytes: 20.36%, Astrocyte: 13.97%, Microglia: 5.12%, Endothelial: 0.63%. AD: Clinical diagnosis of Alzheimer's dementia, MCI: Mild cognitive impairment, NCI: No cognitive impairment.

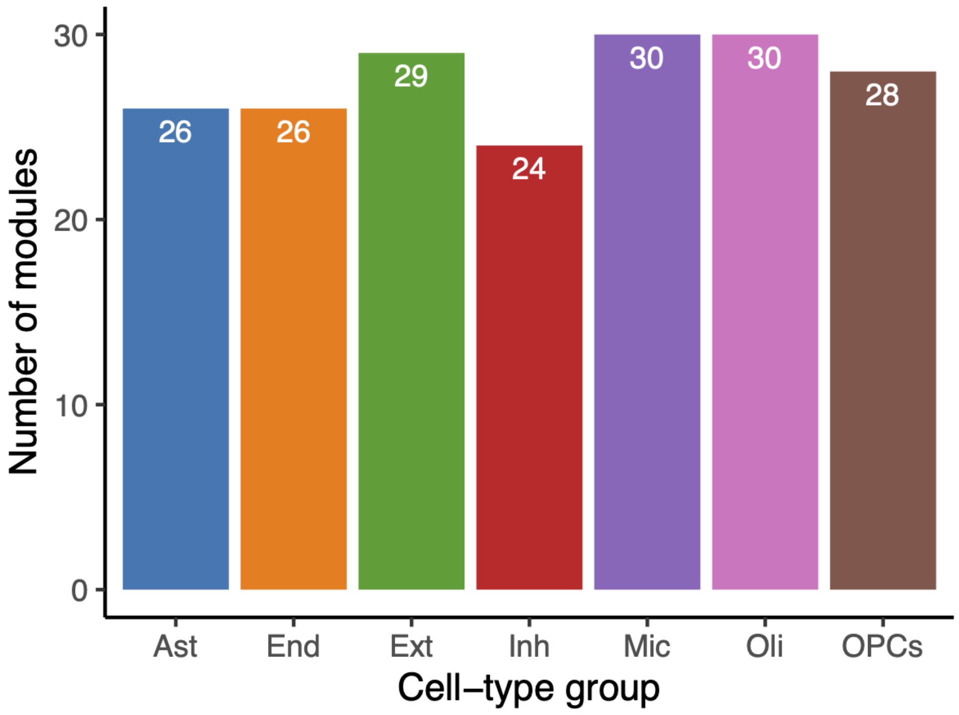

**Supplementary Figure 3: Number of modules by network.**
Ast: Astrocytes, end: Endothelial cells, ext: Excitatory neurons, Inh: inhibitory neurons, mic: Microglia, oli: Oligodendrocytes, and opc: Oligodendrocyte precursor cells.

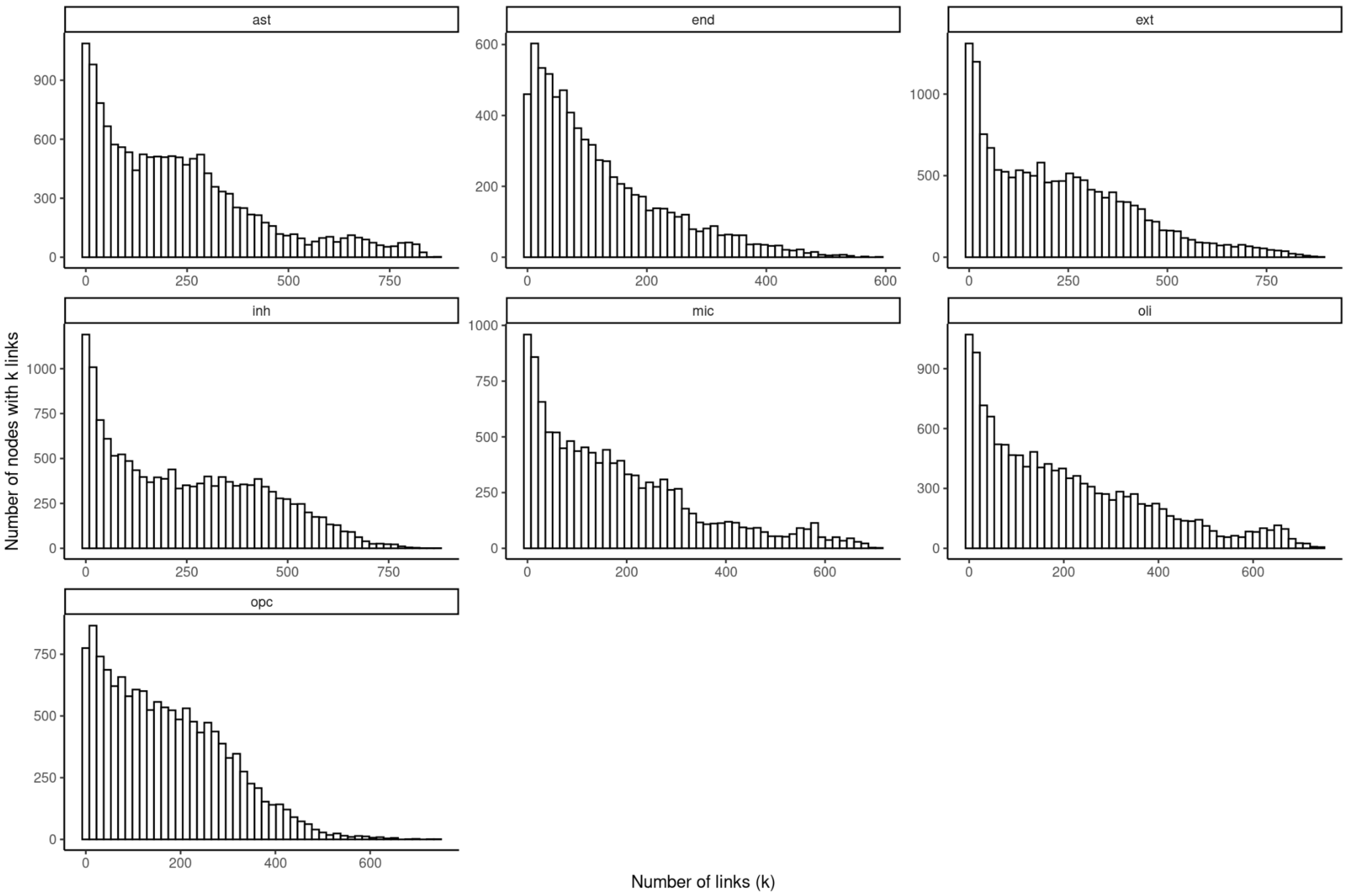

**Supplementary Figure 4: The single-nucleus coexpression networks follow a scale-free topology.** X axis for the number of links (connections) and Y axis for the number of nodes, for each network. Ast: Astrocytes, end: Endothelial cells, ext: Excitatory neurons, Inh: inhibitory neurons, mic: Microglia, oli: Oligodendrocytes, and opc: Oligodendrocyte precursor cells.

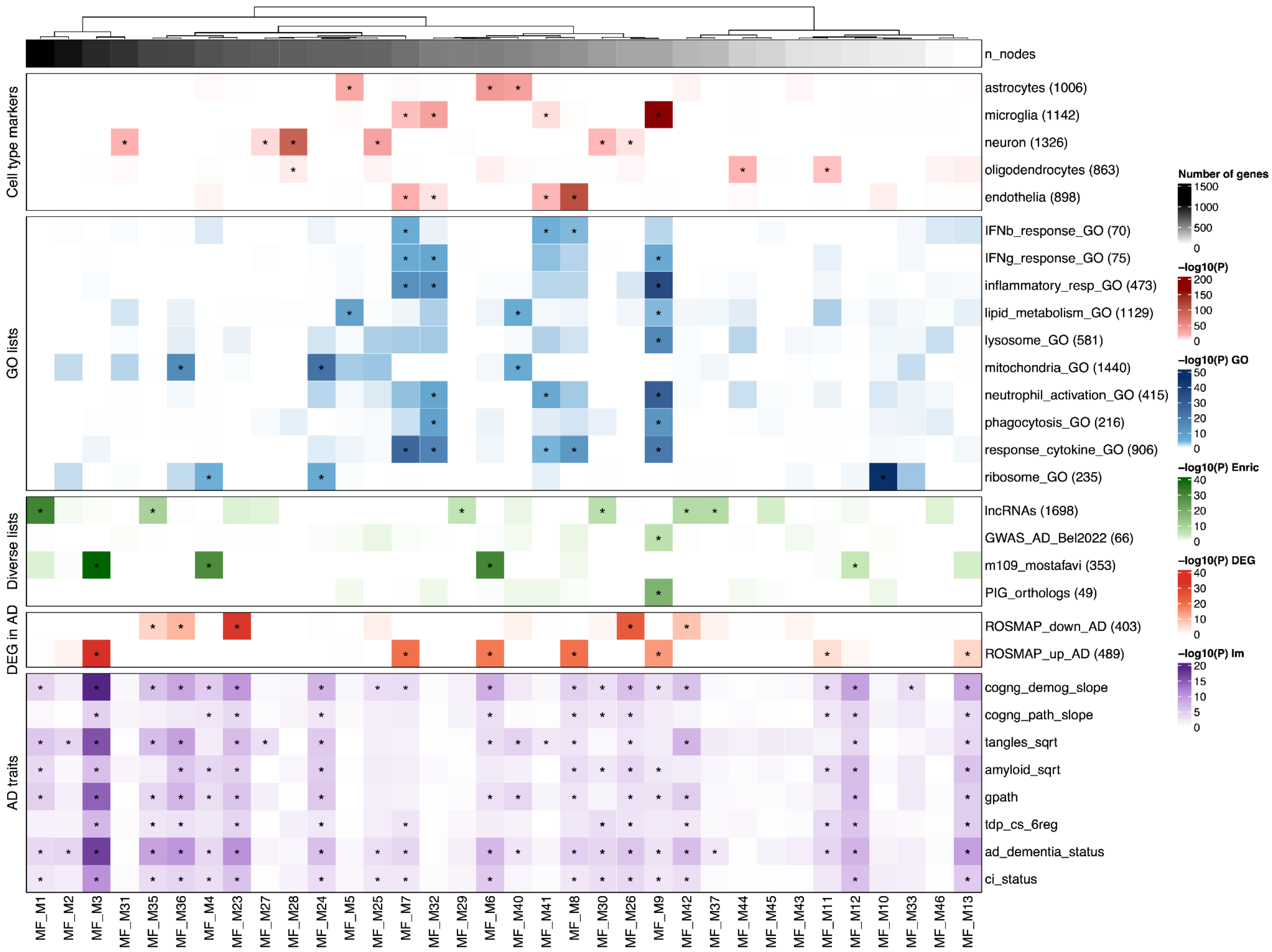

**Supplementary Figure 5: The coexpression modules of the dorsolateral prefrontal cortex (DLPFC) bulk RNASeq (n=1,210 individuals).**

a) First heatmap orders the modules by size. b-e) Fisher exact test results between gene lists and the module average expression. The lists include cell marker genes from Johnson et al. 2022 (Johnson et al. 2022); gene ontology (GO) terms (Ashburner et al. 2000; Gene Ontology Consortium et al. 2023); lncRNAs (Gencode 37); AD GWAS from Bellenguez et al. 2022 (Bellenguez et al. 2022); module 109 previously published by Mostafavi et al., 2018 (Mostafavi et al. 2018); Plaque-induced gene list from Chen et al., 2022 (Chen et al. 2020); genes down and up-regulated in AD from the bulk dataset with 1,210 individuals. The *P*-values were adjusted by computing multiple testing corrections, and asterisks highlight modules with Bonferroni *P*-value < 0.05. f) Association results between the module's average expression and AD traits. The *P*-values are from the linear or logistic regression after adjustment by age, sex, and years of education. Asterisks indicate that the results reached the significance threshold (Bonferroni *P*-value < 0.05).

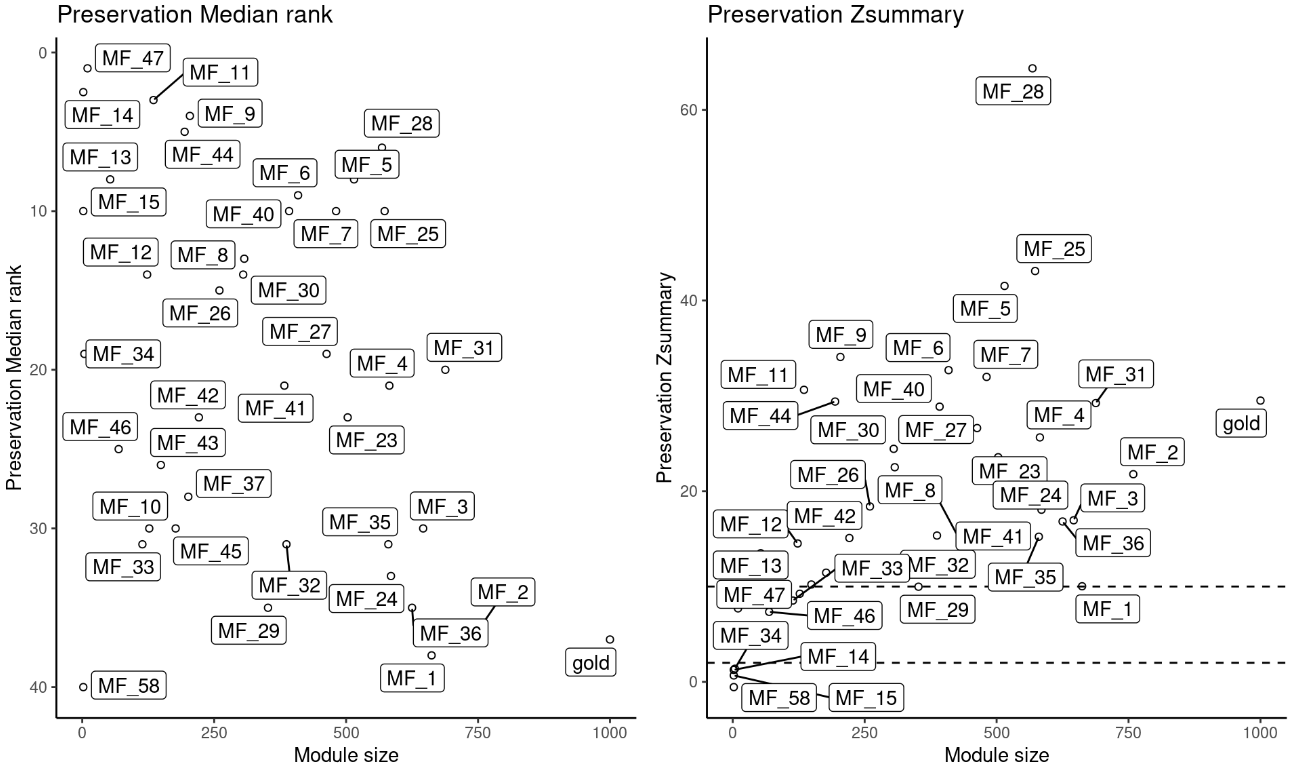

**Supplementary Figure 6: Module preservation between the updated DLPFC (n=1,210 individuals) and Mostafavi et al. 2018 modules.**

a) The composite statistic medianRank (y-axis) as a function of the module size. Each dot is one module labeled accordingly. b) The Zsummary statistic equation (y-axis) as a function of the module size (x-axis). Dashed lines represent the cut-off for the Kruskal–Wallis test for 2 and 10 where modules > 2 are considered preserved or > 10 highly preserved. Gold is a random module.

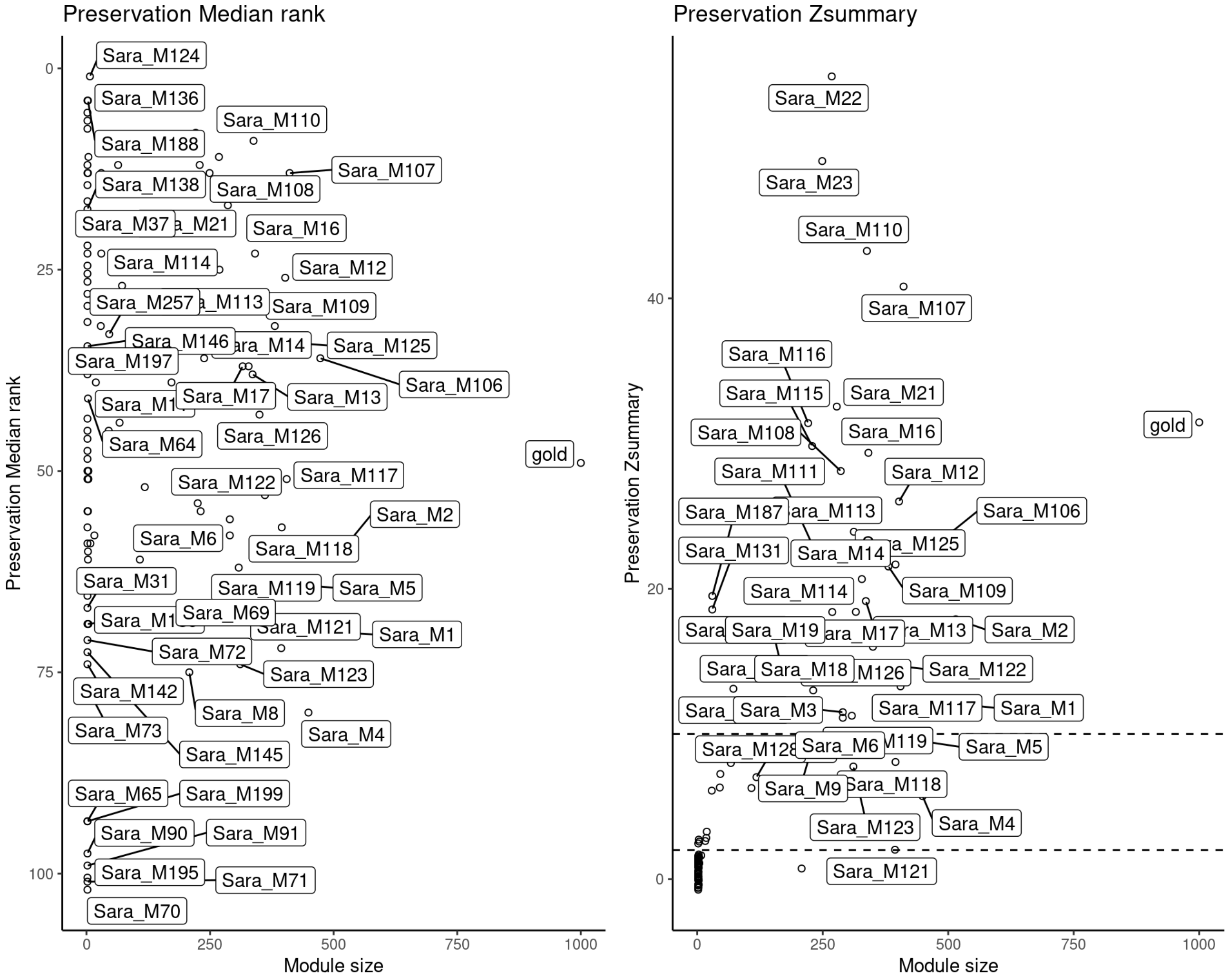

**Supplementary Figure 7: Are the modules from Mostafavi et al. 2018 preserved in the updated DLPFC (n=1,210 individuals) network?**

a) The composite statistic medianRank (y-axis) as a function of the module size. Each dot is one module labeled accordingly. b) The Zsummary statistic equation (y-axis) as a function of the module size (x-axis). Each dot is one module labeled accordingly. Dashed lines represent the cut-off for the Kruskal–Wallis test for 2 and 10 where modules > 2 are considered preserved or > 10 highly preserved. Gold is a random module.

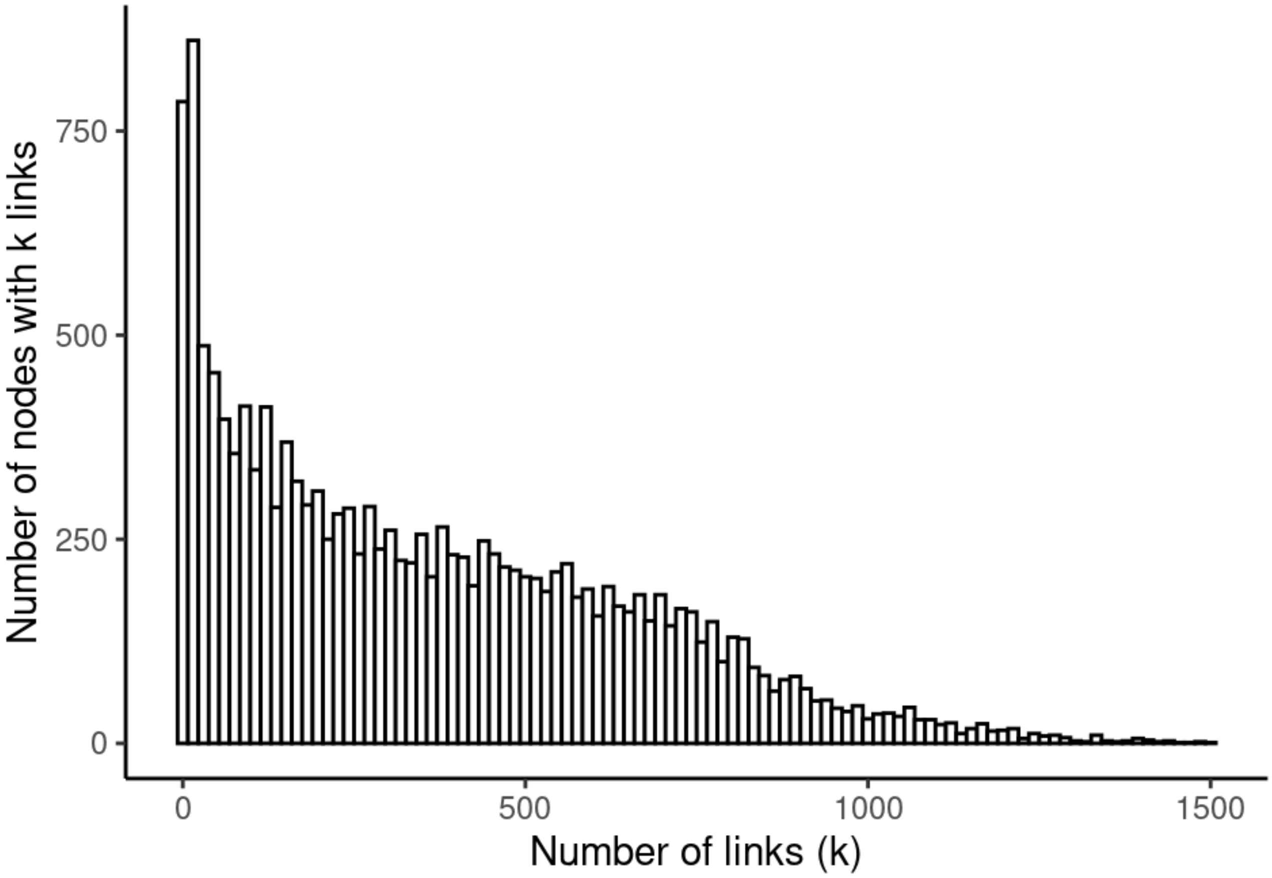

**Supplementary Figure 8: DLPFC network distribution.**
The number of nodes (y-axis) by the number of connections (x-axis).

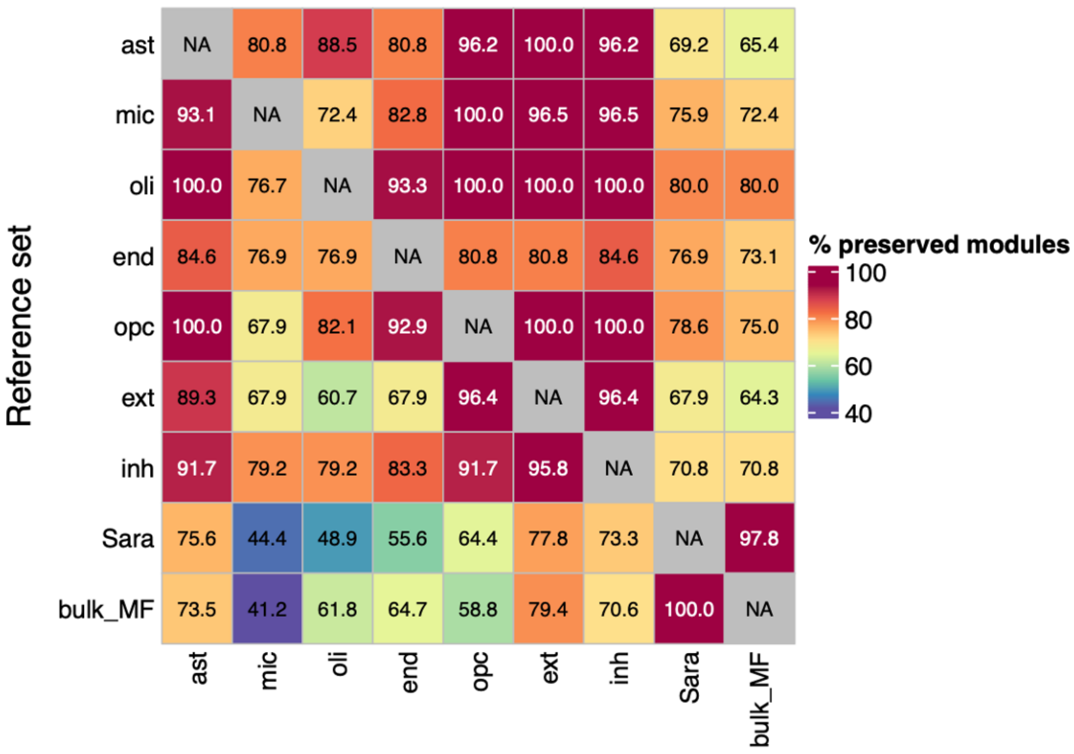

**Supplementary Figure 9: Heatmap showing the percentage of modules preserved across the coexpression networks.**

Heatmap showing the percentage of modules preserved across the coexpression networks according to Zsummary statistic equation based on density and connectivity. Red colors indicate high preservation while purple indicate low preservation values. The (Sara) bulk RNASeq networks are from the previously published Mostafavi et al. *Nat Neuroscience*, 2018 and the updated one (bulk_MF) from this study which included 1,210 individuals.

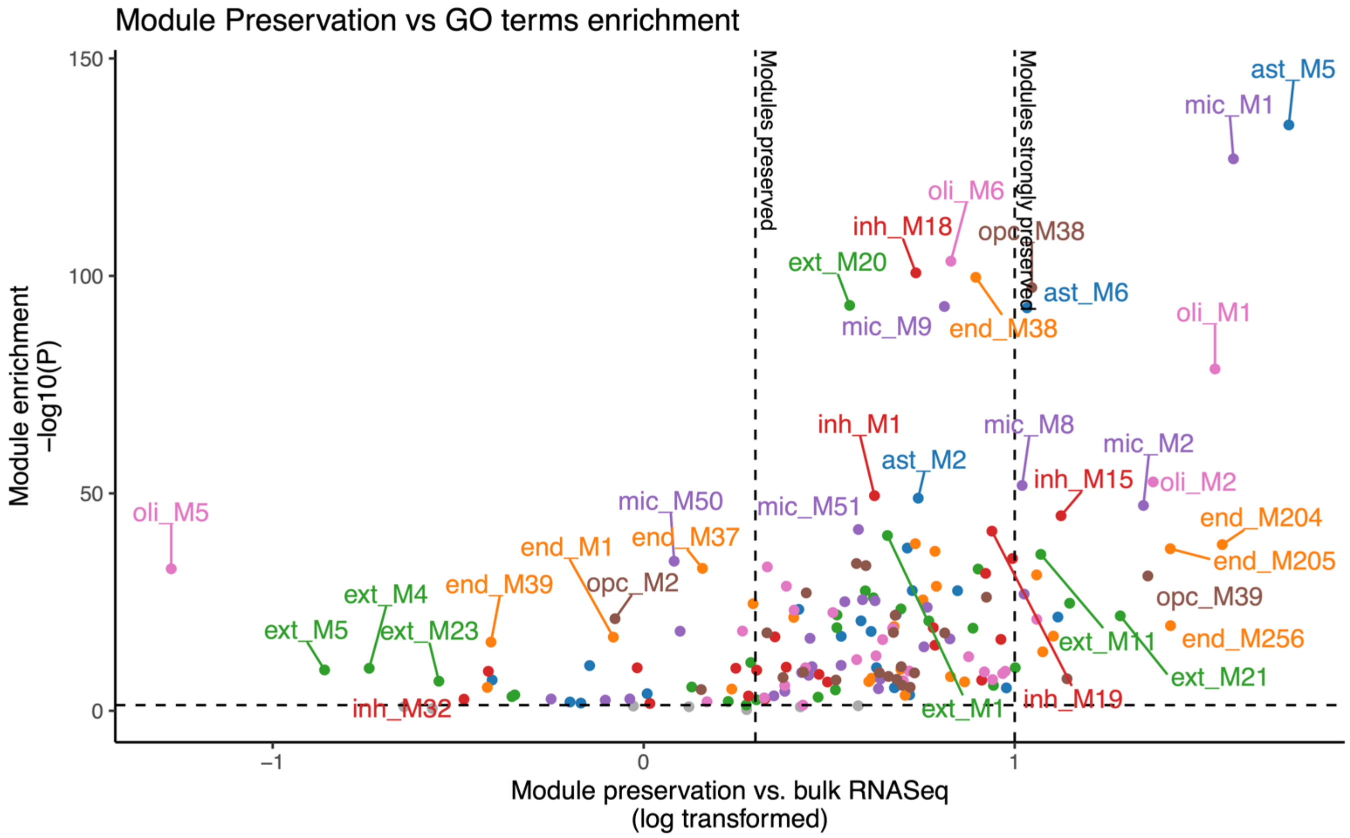

**Supplementary Figure 10: Comparison between module preservation against bulk RNASeq and functional enrichment.**

Module preservation is shown in the x-axis (as $\log_{10}$(Zsummary)) while the y-axis shows the functional enrichment analysis for Gene ontology terms (best term $-\log_{10}$($P$-value). Vertical dashed lines indicate module preservation thresholds: not preserved < 2, moderately preserved if ≥ 2 and < 10, and highly preserved if Zsummary ≥ 10. Horizontal dashed lines indicate $P$-value threshold of 0.05 for terms from Gene ontology.

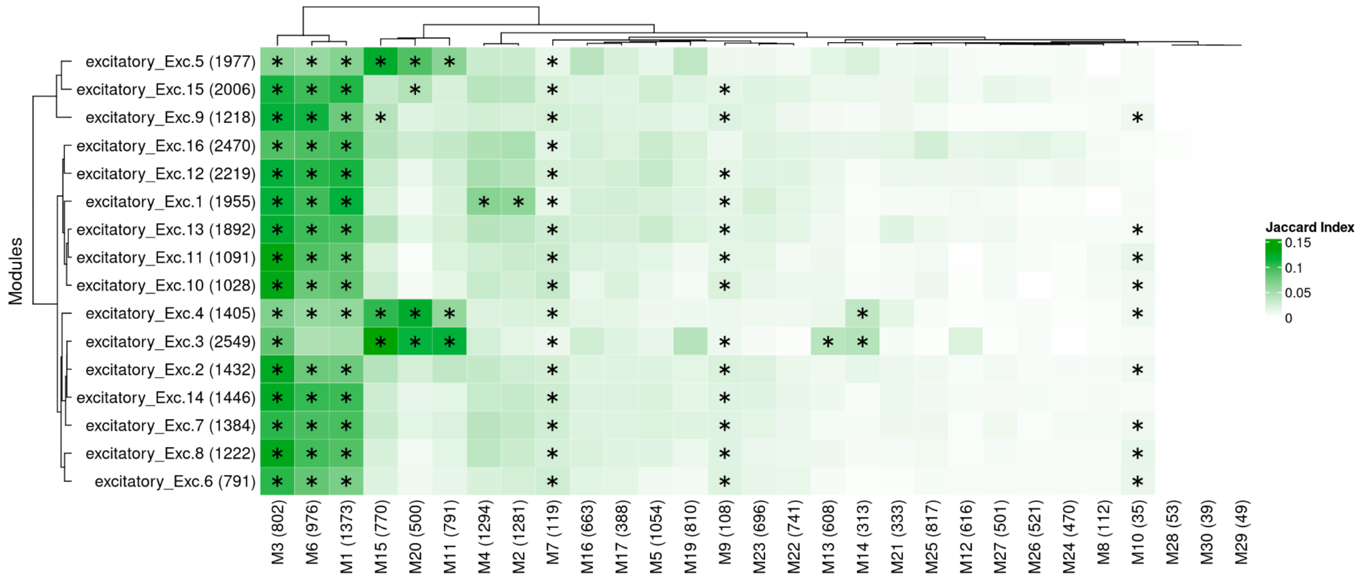

**Supplementary Figure 11: Do the excitatory neurons modules recapitulate sub-cell type clusters or states?**

Fisher exact test results for enrichment analysis between the excitatory neuron modules, and the sub-cell types of marker genes (Green et al. 2024). Asterisks indicate if the results reached a significance threshold, Bonferroni *P*-value < 0.05. A stronger color indicates higher Jaccard index values.

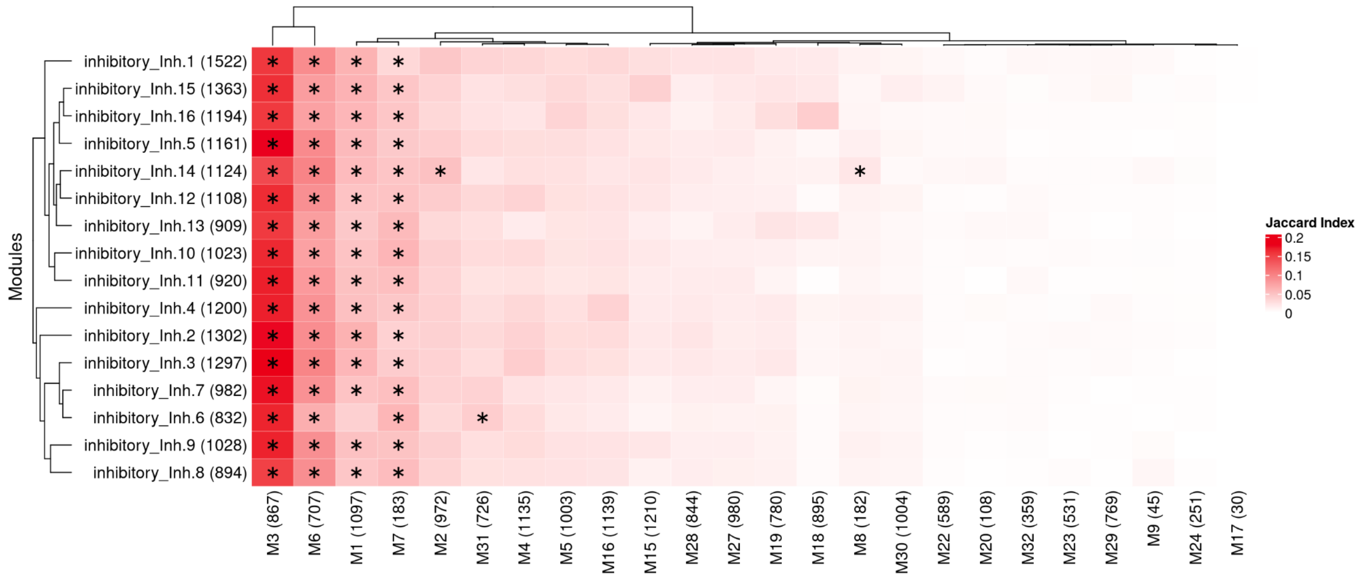

**Supplementary Figure 12: Do the inhibitory neurons modules recapitulate sub-cell type clusters or states?**

Fisher exact test results for enrichment analysis between the inhibitory neuron modules, and the sub-cell types of marker genes (Green et al. 2024). Asterisks indicate if the results reached a significance threshold, Bonferroni *P*-value < 0.05. A stronger color indicates higher Jaccard index values.

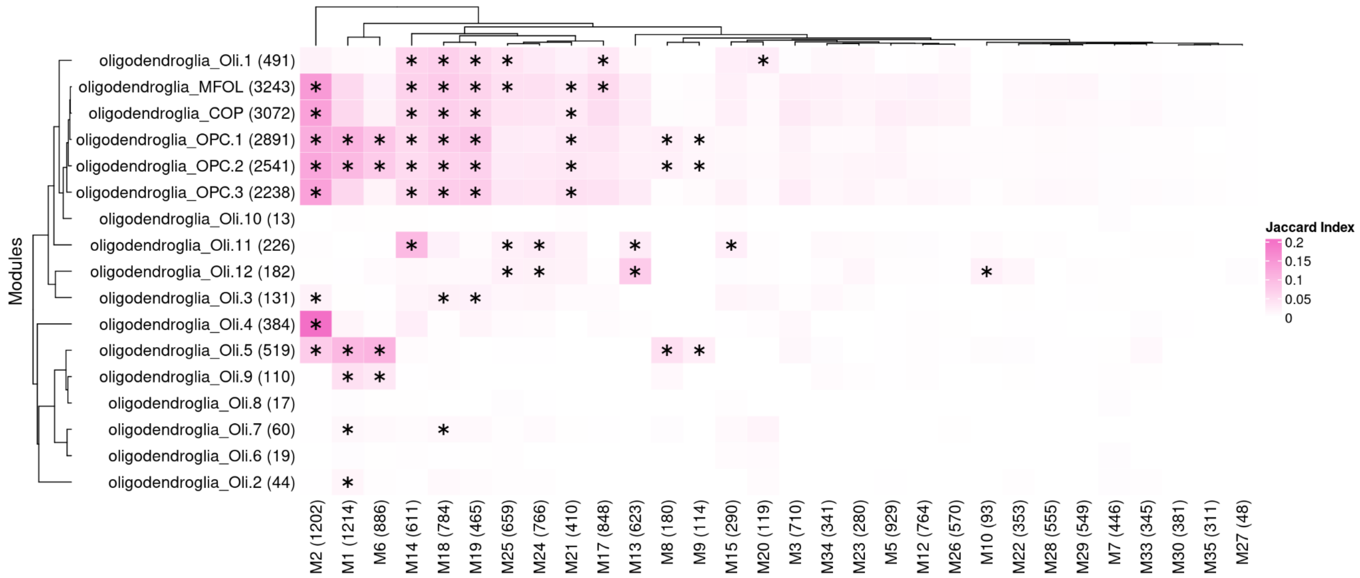

**Supplementary Figure 13: Do the oligodendrocytes modules recapitulate sub-cell type clusters or states?**

Fisher exact test results for enrichment analysis between the oligodendrocytes modules, and the sub-cell types of marker genes (Green et al. 2024). Asterisks indicate if the results reached a significance threshold, Bonferroni *P*-value < 0.05. A stronger color indicates higher Jaccard index values.

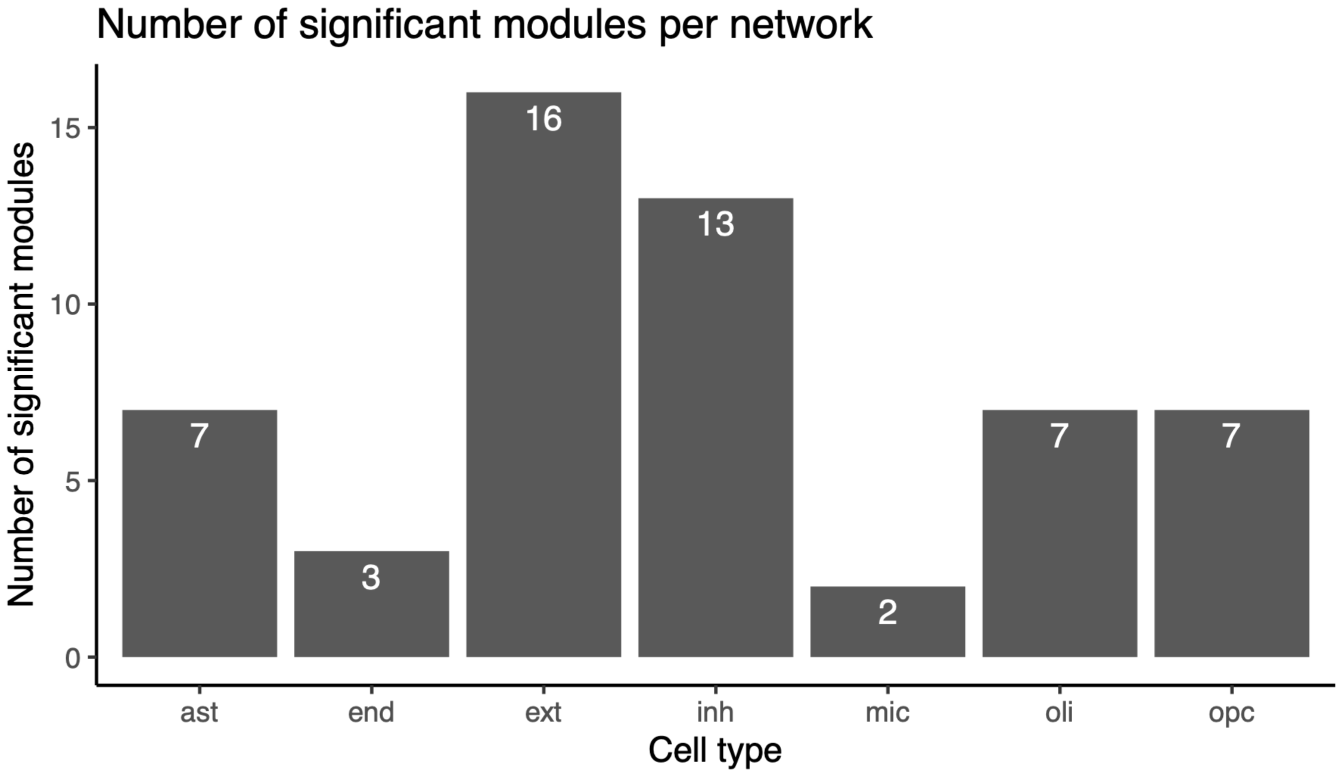

**Supplementary Figure 14: Number of significant modules per network.**
Barplot showing the summarized results from the regression analysis of the module's average expression with eight AD-related traits. FDR *P*-value < 0.05 for significance threshold. Ast: Astrocytes, end: Endothelial cells, ext: Excitatory neurons, Inh: inhibitory neurons, mic: Microglia, oli: Oligodendrocytes, and opc: Oligodendrocyte precursor cells.

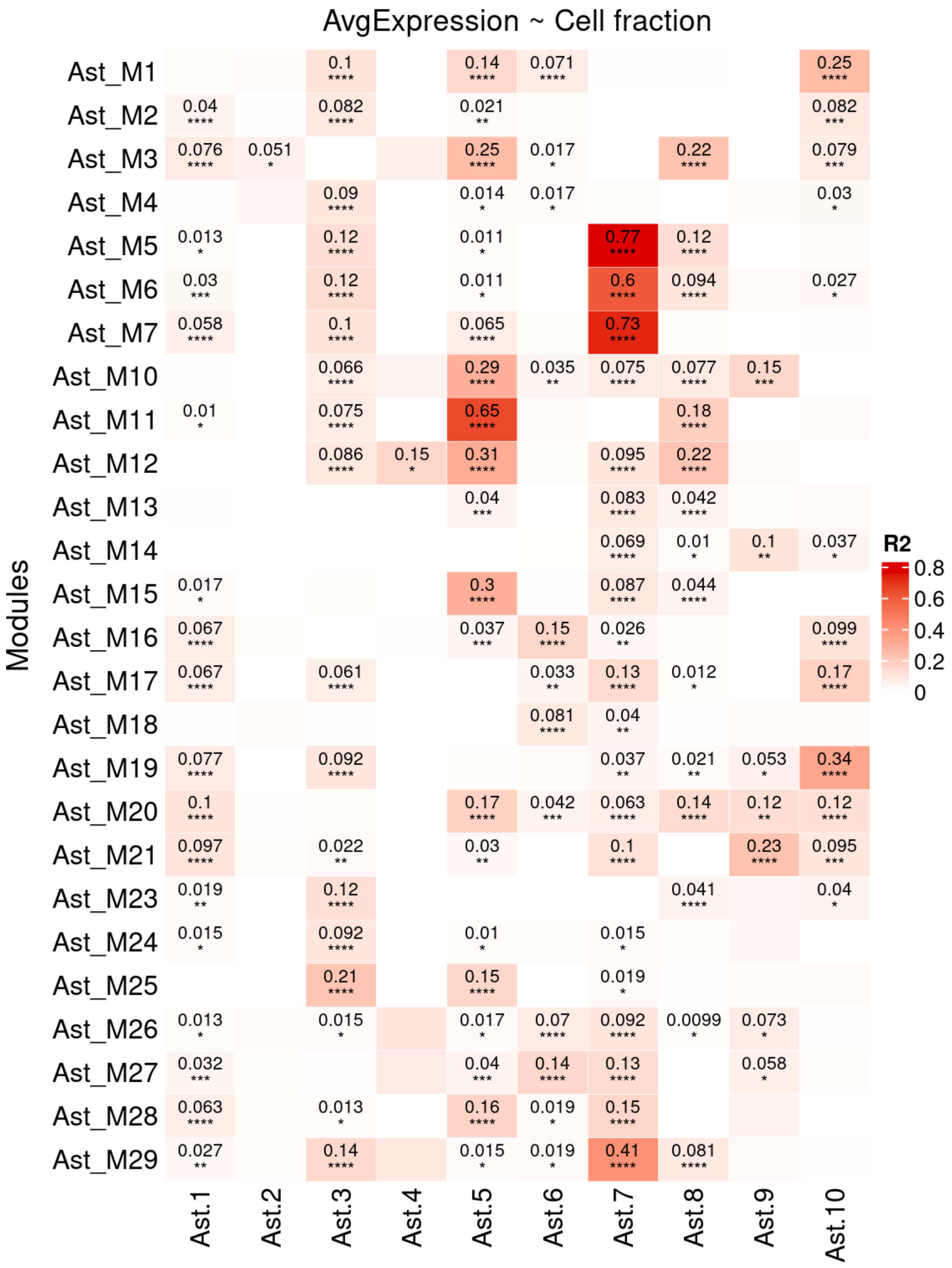

**Supplementary Figure 15: Linear regression between the module's average expression and astrocyte cell fractions.**

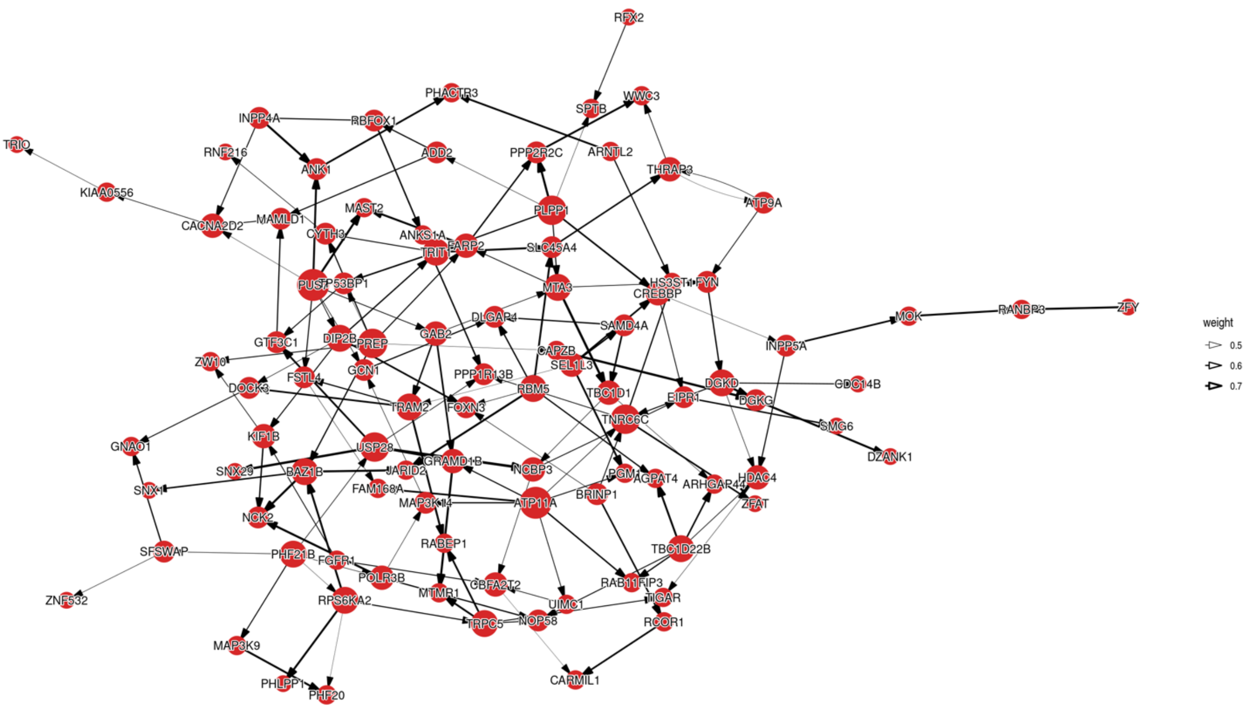

**Supplementary Figure 16: The intra-module Bayesian network for the top 100 genes in the inh_M6.** Top genes were selected according to their association with AD pathology and cognitive decline measurements. The arrows indicate the direction of causality effect, and edge thickness is related to the weights after 5,000 interactions. Node size is proportional to the node's connectivity.
